# Molecular characterization of projection neuron subtypes in the mouse olfactory bulb

**DOI:** 10.1101/2020.11.30.405571

**Authors:** Sara Zeppilli, Tobias Ackels, Robin Attey, Nell Klimpert, Kimberly D. Ritola, Stefan Boeing, Anton Crombach, Andreas T. Schaefer, Alexander Fleischmann

## Abstract

Projection neurons (PNs) in the mammalian olfactory bulb (OB) receive direct input from the nose and project to diverse cortical and subcortical areas. Morphological and physiological studies have highlighted functional heterogeneity, yet no molecular markers have been described that delineate PN subtypes. Here, we used viral injections into olfactory cortex and fluorescent nucleus sorting to enrich PNs for high-throughput single nucleus and bulk RNA deep sequencing. Transcriptome analysis and RNA *in situ* hybridization identified three mitral and five tufted cell populations with characteristic transcription factor network topology and cell adhesion and excitability-related gene expression. Finally, by integrating bulk and snRNA-seq data we propose that different mitral cell populations selectively project to different regions of olfactory cortex. Together, we have identified potential molecular and gene regulatory mechanisms underlying PN diversity and provide new molecular entry points into studying the diverse functional roles of mitral and tufted cell subtypes.

## Introduction

The mammalian olfactory system is unique among sensory systems in that it bypasses the thalamus: olfactory receptor neurons (ORNs) in the nose project to the olfactory bulb (OB), a forebrain structure containing – in the mouse – approximately 500,000 neurons per hemisphere (Parrish-Aungst et al., 2007). There, they synapse onto various interneurons and projection neurons. The latter directly project to a variety of cortical structures, including the anterior olfactory nucleus, piriform cortex, cortical amygdala and the lateral entorhinal cortex (Ghosh et al., 2011; Haberly and Price, 1977; Miyamichi et al., 2011; Sosulski et al., 2011). This places OB projection neurons at a pivotal position to distribute processed olfactory information broadly across the brain.

Each ORN in the mouse expresses only one of approximately 1000 olfactory receptor genes (Buck and Axel, 1991; Niimura, 2012; Zhang and Firestein, 2002). ORNs expressing the same receptor project axons onto defined spherical structures, glomeruli (Mori and Sakano, 2011), containing a variety of neuropil including the apical dendrites of 10-50 projection neurons (Bartel et al., 2015; Schwarz et al., 2018). Historically, OB projection neurons have been divided into mitral and tufted cells (MCs, TCs), largely based on their soma location and dendritic and axonal projection pattern (**Figure 1-figure supplement 1**) (Haberly and Price, 1977; Imamura et al., 2020; Mori et al., 1983; Orona et al., 1984): MC somata are located predominantly in a thin layer with their dendrites covering the deeper part of the OB external plexiform layer. Their axons project to a wide range of structures including posterior piriform cortex. TC axons, on the other hand, are restricted to more anterior forebrain structures and their cell bodies are distributed across the external plexiform layer, with dendrites largely restricted to superficial layers. Within the TC population, several subdivisions have been made into deep, middle, superficial and external TCs, largely based on soma position. MCs on the other hand are often morphologically described as a largely homogeneous population. However, branching patterns of lateral dendrites as well as soma size and apical dendrite length might allow for further subdivision (Mouradian and Scott, 1988; Orona et al., 1984; Schwarz et al., 2018). Moreover, projection patterns might differ based on soma position along the dorsomedial-ventrolateral axis of the OB (Inokuchi et al., 2017).

In parallel to this morphological diversity, numerous studies have described physiological heterogeneity both as a result of differential inputs from granule cells onto TCs and MCs (Christie et al., 2001; Ezeh et al., 1993; Geramita et al., 2016; Phillips et al., 2012) as well as intrinsic excitability and possibly glomerular wiring (Burton and Urban, 2014; Gire et al., 2019). Consequently, TCs respond more readily, with higher peak firing rates, and to lower odor concentration *in vivo* (Griff et al., 2008; Kikuta et al., 2013; Nagayama et al., 2014), and earlier in the respiration cycle compared to MCs (Ackels et al., 2020; Fukunaga et al., 2012; Igarashi et al., 2012; Jordan et al., 2018; Phillips et al., 2012).

Within the TC and MC populations, biophysical heterogeneity has been more difficult to tie to specific cell types or subtypes. MCs show diversity in biophysical properties that is thought to aid efficient encoding of stimulus-specific information and is, at least in part, experience-dependent (Angelo et al., 2012; Burton et al., 2012; Padmanabhan and Urban, 2010; Tripathy et al., 2013). Both *in vivo* and *in vitro* recordings suggest that a subset of MCs show regular firing, whilst others show ‘stuttering’ behavior characterized by irregular action potential clusters (Angelo et al., 2012; Balu et al., 2004; Bathellier et al., 2008; Buonviso et al., 2003; Carey and Wachowiak, 2011; Desmaisons et al., 1999; Fadool et al., 2011; Friedman and Strowbridge, 2000; Margrie and Schaefer, 2003; Padmanabhan and Urban, 2010; Schaefer et al., 2006). While TCs are heterogeneous, with for example external TCs displaying prominent rhythmic bursting, driving the glomerular circuitry into long-lasting depolarizations *in vitro* (De Saint Jan et al., 2009; Gire and Schoppa, 2009; Gire et al., 2019; Najac et al., 2011), a systematic assessment of biophysical variety is lacking so far. Moreover, differential centrifugal input from cortical and subcortical structures might further amplify this overall heterogeneity both between MCs and TCs as well as potentially within those different classes (Boyd et al., 2012; Kapoor et al., 2016; Markopoulos et al., 2012; Niedworok et al., 2012; Otazu et al., 2015).

Thus, anatomical projection patterns, *in vivo* odor responses, and intrinsic properties are known to show substantial variability across different projection neurons. Systematic investigation of different projection neurons, however, has been hampered by a scarcity of specific molecular tools. Interneuron diversity, on the other hand, in general has received considerable attention with numerous studies including in the OB (Parrish-Aungst et al., 2007; Tavakoli et al., 2018) aiming to provide a systematic assessment of morphology, physiology, chemotype and the basis for genetic targeting of distinct types of interneurons. For OB projection neurons, however, only little information about chemotypes is available at this point: *Pcdh1* and *Tbx21* (Haddad et al., 2013; Nagai et al., 2005) have been shown to be selectively expressed in a subset of OB projection neurons. CCK distinguishes a subset of TCs (superficial TCs, (Liu and Shipley, 1994; Seroogy et al., 1985; Short and Wachowiak, 2019; Sun et al., 2020). Vasopressin expressing cells might constitute a further subset of superficial TCs (Lukas et al., 2019), and recently the *Lbhd2* gene has been used to obtain more specific genetic access to MCs (Koldaeva et al., 2020). Heterogeneous expression of both the GABAa receptor as well as voltage-gated potassium channel subunits have been observed (Padmanabhan and Urban, 2010; Panzanelli et al., 2005), albeit not linked to specific cell types. Expression of axon guidance molecules such as *Nrp2* might further allow subdivision of projection neurons across the OB (Inokuchi et al., 2017).

Hence, while some molecular markers can be used to define specific subsets of projection neurons, this description is far from complete. A comprehensive molecular definition of projection neuron types would help to classify and collate existing biophysical, morphological and physiological data and delineate the distinct output streams of the OB. Moreover, it would provide a platform upon which further focused experimental approaches could be tied.

Single cell or single nucleus RNA sequencing has been used effectively to map out cell type across a variety of brain areas (Macosko et al., 2015; Zeisel et al., 2018), including inhibitory interneurons in the mouse OB (Tepe et al., 2018). As M/TCs constitute only ∼10% of all OB neurons, we decided to enrich for projection neurons for single nucleus (sn)RNA-seq. We then combined snRNA-seq with bulk RNA deep seq for OB neurons projecting to different cortical areas, thereby allowing us to disentangle different projection neuron classes by target area. We found that indeed both MCs and TCs fall into several, separable types, defined by expression of both common and overlapping gene regulatory networks. This work will therefore provide a molecular entry point into disentangling the diversity of OB projection neurons and defining the functional roles of different MC/TC types.

## Results

### Single nucleus RNA sequencing of olfactory bulb projection neurons distinguishes mitral and tufted cell types

To characterize the molecular diversity of OB projection neurons, we devised two complementary experimental strategies. We used viral targeting and Fluorescence-Activated Nuclei Sorting (FANS) to enrich for OB projection neurons, and we then characterized their transcriptomes using single nucleus RNA sequencing (snRNA-seq) and bulk RNA deep sequencing **(Figure 1A, Figure 1-figure supplement 2)**.

**Figure 1:**
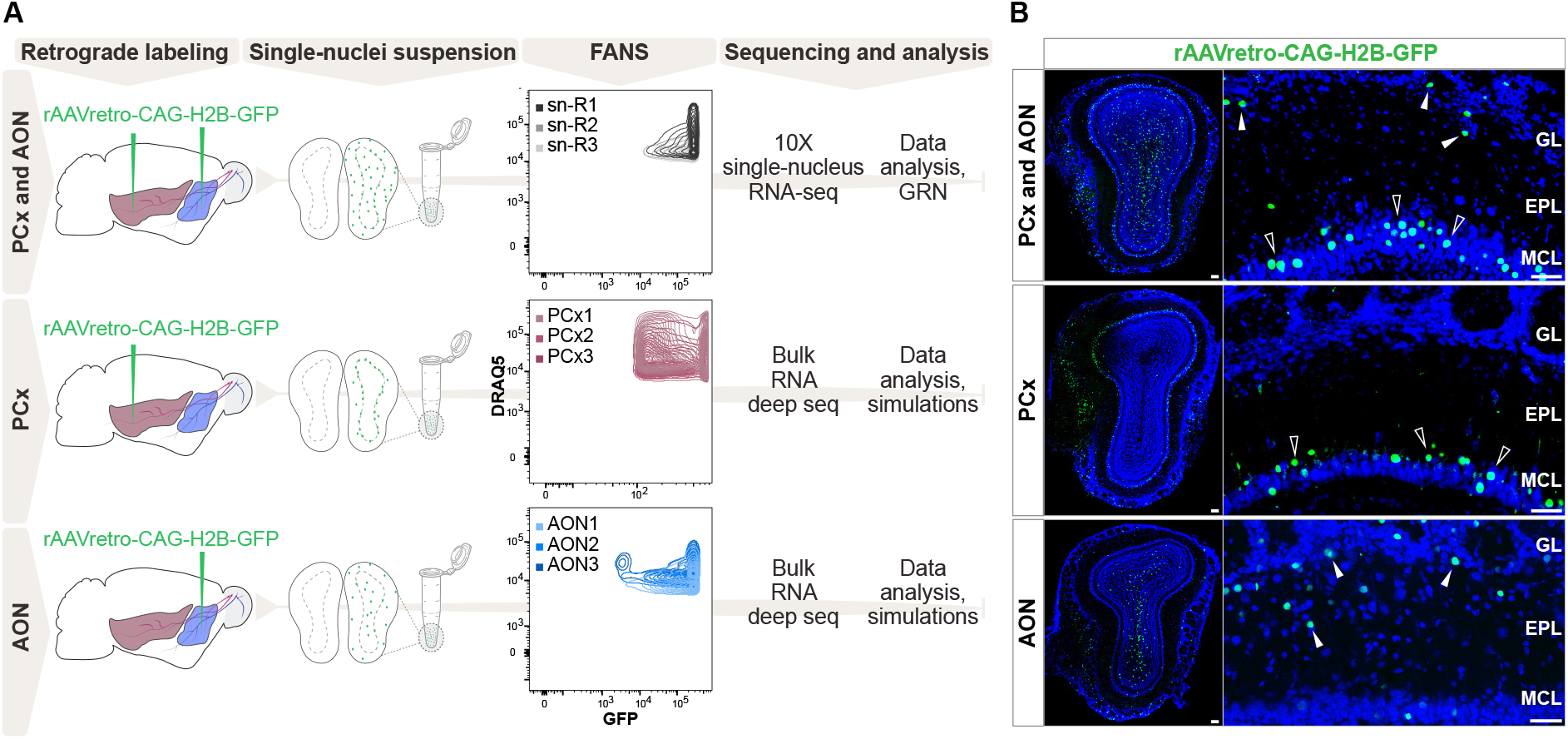
Comprehensive molecular profiling of olfactory bulb projection neurons. **(A)** Schematic representation of experimental design. Top: after injection of rAAVretro-CAG-H2B-GFP into PCx and AON, single nuclei were dissociated from 3 mice (single nuclei (sn) R1,2,3: replicates 1,2,3) and sorted using Fluorescence-activated Nuclei Sorting (FANS). The population of nuclei is selected based on GFP and DRAQ5 (far-red fluorescent DNA dye). See **Figure 1-figure supplement 2** for detailed FANS plots. Sorted nuclei were sequenced using 10X single-nucleus RNA-seq. Middle and bottom: after injection of rAAVretro-CAG-H2B-GFP into PCx (middle) or AON (bottom), single nuclei were dissociated from 3 mice for each injection site and sorted using FANS (as described above and **Figure 1-figure supplement 2**). RNA extracted from sorted nuclei was sequenced using bulk RNA deep sequencing. PCx: Piriform Cortex; AON: Anterior Olfactory Nucleus; R: replicate; GRN: Gene Regulatory Network. **(B)** Representative coronal sections and high magnification images showing GFP expression (in green) in the olfactory bulb after injection of rAAVretro-CAG-H2B-GFP into PCx and AON (top), PCx only (middle), and AON only (bottom). Injection of the virus into PCx and AON resulted in GFP-expressing nuclei located in the mitral cell (empty arrowheads), external plexiform, glomerular (white arrowheads), and granule cell layers; injection into PCx resulted in GFP-expressing nuclei located in the mitral cell layer (empty arrowheads); injection into AON resulted in GFP-expressing nuclei located in the external plexiform and glomerular layers (white arrowheads) and granule cell layers. GL: glomerular layer; EPL: external plexiform layer; MCL: mitral cell layer; GCL: granule cell layer. Neurotrace counterstain in blue. Scale bars, 100μm and 50μm (high magnification).

First, we injected a retrogradely transported Adeno-Associated Virus expressing nuclear GFP (rAAV-retro-CAG-H2B-GFP (Tervo et al., 2016)) into multiple sites along the antero-posterior axis of the olfactory cortex (**Figure 1A)**. Histological analysis revealed that virus injections resulted in GFP expression in a heterogeneous population of OB cells labelling cells in the mitral cell, external plexiform, glomerular and granule layers **(Figure 1B)**. Sparse GFP expression in putative periglomerular and granule cells may have resulted from viral infection of migrating immature neurons from the rostral migratory stream or from diffusion of the virus from the injection site.

We dissected the olfactory bulbs of three injected mice, generated three independent replicates of single nuclei, enriched for GFP expression using FANS (**Figure 1-figure supplement 2)**, and performed snRNA-seq using 10X Genomics technology **(Figure 1A)**. We performed a detailed quality check of the individual replicates, then combined nuclei for downstream analyses **(Figure 2-figure supplement 1)**. We analyzed a total of 31,703 nuclei, grouped in 22 clusters that were annotated *post hoc* based on the expression of established marker genes for excitatory and inhibitory neurons and glial cell populations **(Figure 2A-C)**. We initially used the combinatorial expression patterns of glutamatergic markers and previously characterized M/T cell markers (*Vglut1, Vglut2, Vglut3, Tbx21, Pcdh21, Thy1)* to identify putative OB projection neurons, comprising 23.66% (n=7504) of all nuclei.

**Figure 2:**
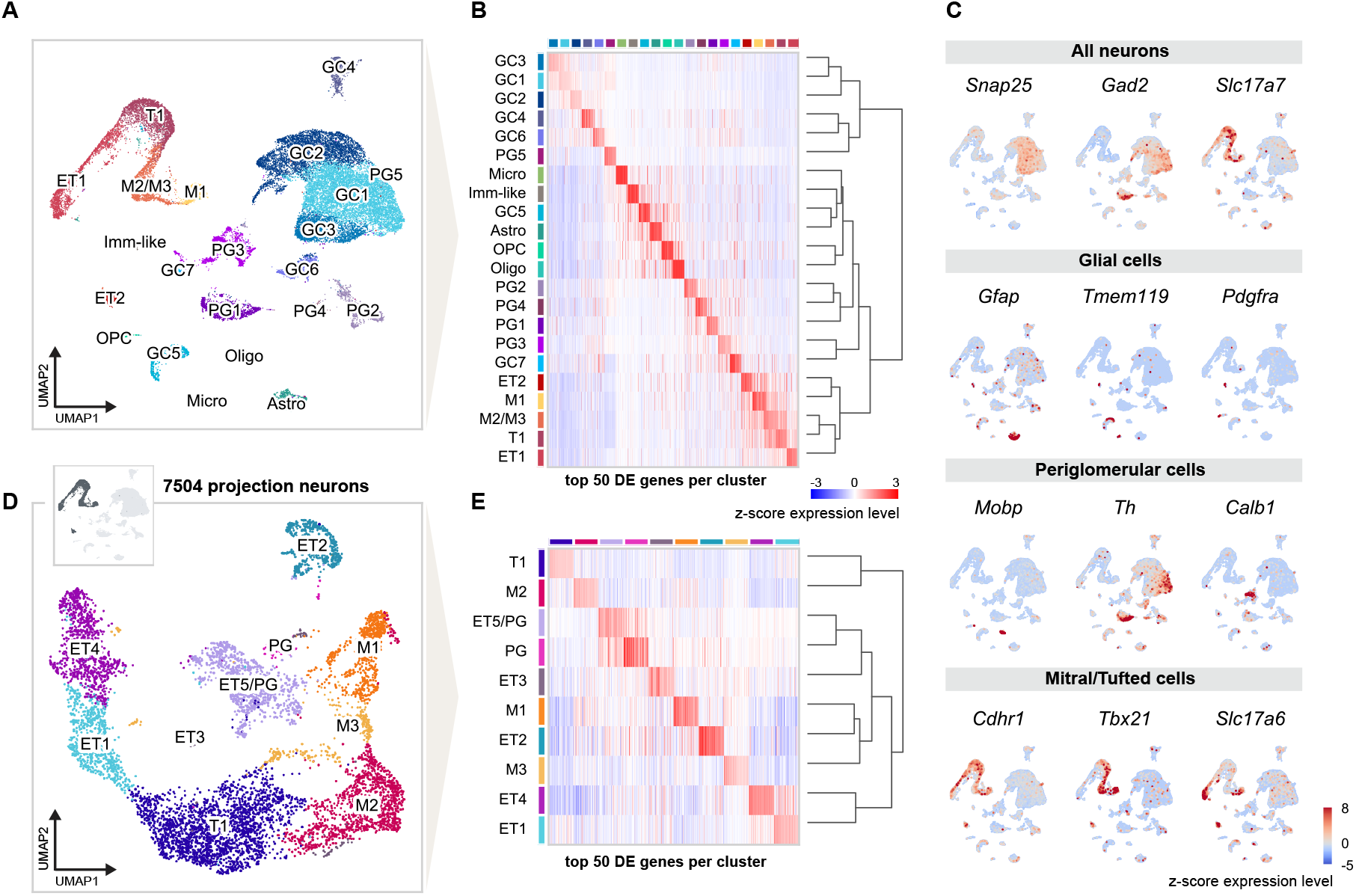
Single nucleus RNA sequencing distinguishes distinct cell types and molecular signatures of OB projection neurons. **(A)** UMAP representation of gene expression profiles of 31,703 single nuclei combined from all replicates (1, 2, 3) of mice injected into both AON and PCx, grouped into 22 clusters color-coded by cell type membership (GC: granule cell, PG: periglomerular cell, OPC: oligodendrocyte precursor cell, Micro: microglia, Astro: astrocyte, Oligo: oligodendrocyte, ET: external tufted cell, M: mitral cell, T: tufted cell, Imm-like: Immature-like cell). See **Figure 2-figure supplement 1** for detailed quality check of each replicate. **(B)** Matrixplot showing the z-score expression level of the top 50 differentially expressed (DE) genes for each cell population ordered by hierarchical relationships between distinct clusters. Each column represents the average expression level of a gene in a given cluster, color-coded by the UMAP cluster membership (from **A)**. The dendrogram depicts the hierarchical relationships and is computed from the PCA representation of the data using Pearson correlation as distance measure and link by complete linkage. **(C)** UMAP representations of known marker genes for main cell populations (*Snap25*: neurons; *Gad2*: GABAergic neurons; *Slc17a7*: glutamatergic neurons; *Gfap, Tmem119, Pdgfra, Mobp*: glial cells; *Th, Calb1*: periglomerular neurons; *Cdhr1, Tbx21, Slc17a6*: mitral/tufted cells). Nuclei are color-coded by the z-score expression level of each transcript. **(D)** UMAP representation of subclustering from initial clusters M1, M2/M3, T1, ET1 and ET2 (cluster IDs from **Figure A**), selected for the expression of known excitatory and mitral/tufted cell markers (shown in **C)**, showing 7,504 putative projection neurons grouped into 10 distinct types. **(E)** Same matrixplot as described in **B** showing the z-score expression level of the top 50 DE genes for each projection neuron type ordered by hierarchical relationships and color-coded by the UMAP subcluster membership (from **D)**.

Next, we further subclustered the selected profiles, resulting in a total of 10 molecularly distinct subpopulations. To assign preliminary labels to each of these cell types, we used marker genes previously employed in functional or single cell RNA-seq studies (Nagayama et al., 2014; Tepe et al., 2018). We also used available RNA *in situ* hybridization data from the Allen Institute for Brain Science to corroborate our preliminary assignments (**Figure 3-figure supplement 1 and 2**). Our analysis revealed eight molecularly distinct clusters of putative projection neurons and two clusters of putative periglomerular cells **(Figure 2D and E)**. Among projection neurons, we identified three clusters of MCs (M1, M2, M3) and five clusters of middle and external TCs (T1, ET1, ET2, ET3, ET4).

### smFISH validates mitral and tufted cell types

To identify genes selectively expressed in OB projection neurons, we used the R-package glmGamPoi (Ahlmann-Eltze and Huber, 2020). We calculated the combined average raw expression of the top differentially expressed genes for each cell type and found it to be highly specific for each cluster (**Figure 3A and C)**. We then selected a few specific marker genes **(Figure 3B and D)** to validate projection neuron type identity by combining single molecule Fluorescent *In Situ* Hybridization (smFISH) with GFP staining upon rAAV-retro-CAG-H2B-GFP injection in olfactory cortex.

**Figure 3:**
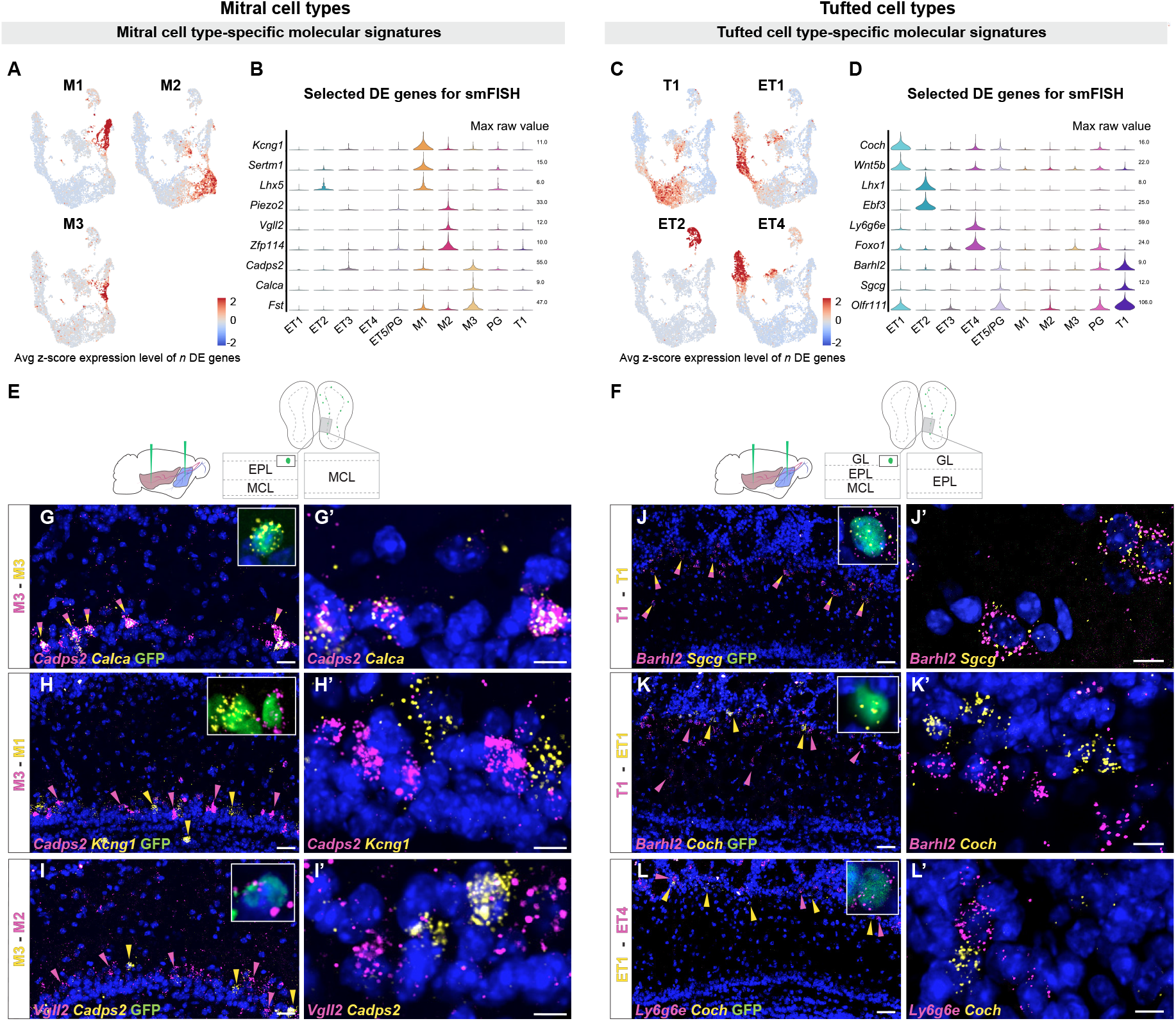
Histological validation of molecularly distinct mitral and tufted cell types. **(A)** Combined average (avg) z-score expression level of the top *n* differentially expressed (DE) genes for each mitral cell type (M1 *n*=14, M2 *n*=11, M3 *n*=10), overlaid on the projection neuron UMAP space (**Figure 2D**). DE genes were selected if their log fold change was greater than 4 (see Methods for details). M1-specific genes: *Kcng1, Lhx1, Sertm1, Gabra2, Doc2b, Cntn6, Olfr1259, Nrp2, C1ql1, Ebf1, Baiap3, Adgrl2, Dsc2, Chrna5;* M2-specific genes: *Piezo2, Vgll2, Zfp114, Nts, Ros1, Samsn1, Grid2, Smpx, Itga4, Itga9, Sema6d;* M3-specific genes: *Cadps2, Calca, Fst, Ets1, Ednra, Cdkn1c, Mustn1, Smoc2, Cnr1, Ccno*. **(B)** Violin plots showing maximum raw expression value of selected mitral cell type-specific DE genes across mitral and tufted cell clusters for further validation with smFISH. **(C)** Combined average (avg) z-score expression level of the top *n* DE genes for each tufted cell type (T1 *n*=9, ET1 *n*=7, ET2 *n*=9, ET4 *n*=6), overlaid on the projection neuron UMAP space (**Figure 2D**). T1-specific genes: *Barhl2, Sgcg, Vdr, Olfr111, Olfr110, Cacna1g, Fam84b, Kcna10, Tspan10;* ET1-specific genes: *Coch, Wnt5b, Rorb, Chst9, Tpbgl, Clcf1, Rxfp1;* ET2-specific genes: *Lhx1, Ebf3, Trp73, Edn1, Ebf2, Nr2f2, Uncx, Psrc1, Dsp;* ET4-specific genes: *Ly6g6e, Foxo1, Siah3, Galnt12, Itga8, Ets2, Grik4*. **(D)** Violin plots showing maximum raw expression value of selected tufted cell type-specific DE genes across mitral and tufted cell clusters for further validation with smFISH. **(E, F)** Schematic representations of the smFISH images for validating projection neuron type-specific selected marker genes upon rAAVretro-CAG-H1B-GFP injection into PCx and AON. The schemes depict the laminar location visualized in the histological images from a coronal section of the ipsilateral hemisphere to the injection site. EPL: external plexiform layer; MCL: mitral cell layer; GL: glomerular layer. **(G - I)** smFISH showing combinatorial expression of mitral cell type-specific marker genes for M1, M2 and M3 cells in the mitral cell layer. High magnifications (top right) show co-labeling of viral GFP with the *in situ* mRNA probe. **(G and G’)** The M3 markers *Cadps2* and *Calca* are co-expressed in subpopulations of cells in the mitral cell layer, indicated by the yellow/magenta arrowheads. **(H and H’)** The M3 marker *Cadps2* and M1 marker *Kcng1* are mutually exclusive in subpopulations of cells in the mitral cell layer, indicated by the magenta and yellow arrowheads respectively. **(I and I’)** The M3 marker *Cadps2* and M2 marker *Vgll2* are mutually exclusive in subpopulations of cells in the mitral cell layer, indicated by the yellow and magenta arrowheads respectively. For additional histological analysis see **Figure 3-figure supplement Figure 1**. **(J - L)** smFISH images showing combinatorial expression patterns of tufted cell type-specific marker genes for validating T1, ET1, ET2 and ET4 clusters as distinct projection neuron types in the external plexiform and glomerular layers. High magnifications (top right) show co-labeling of viral GFP with the *in situ* mRNA probe. As described for the mitral cell types, yellow or magenta arrowheads show mutually exclusive patterns (**K, K’:** T1-ET1 and **L, L’:** ET1-ET4), and yellow/magenta arrowheads show co-expression patterns (**J, J’:** T1-T1). For additional histological analysis see **Figure 3-figure supplement 1**. DAPI counterstain in blue. Scale bars, 50μm and 10μm (high magnifications).

We first investigated MC type-specific gene expression. Differential expression (DE) analysis identified the voltage-gated potassium channel *Kcng1*, the transcriptional regulator LIM homeobox 5 (*Lhx5)* and the serine-rich transmembrane domain 1 (*Sertm1)* as putative M1-specific marker genes. Two-color smFISH revealed selective co-localization of *Kcng1* and *Lhx5* transcripts within the same subpopulation of cells in the MC layer (**Figure 3-figure supplement 1E)**. Furthermore, *Kcng1, Lhx5* and *Sertm1* expression was consistently observed in neurons expressing GFP **(Figure 3H, Figure 3-figure supplement 1B-D)**. Next, DE analysis identified the mechanosensory ion channel *Piezo2*, the transcription cofactor vestigial like family member 2 (*Vgll2)* and the zinc finger protein 114 (*Zfp114)* as putative M2-specific markers. Two-color smFISH revealed extensive co-localization of *Piezo2* and *Vgll2* transcripts within the same subpopulation of cells in the mitral cell layer **(Figure 3-figure supplement 1J)**, and co-localization of M2-specific marker genes with GFP **(Figure 3I, Figure 3-figure supplement 1F and G)**. Finally, we identified the calcium-dependent secretion activator 2 (*Cadps2)*, calcitonin (*Calca)* and follistatin (*Fst)* as putative M3-specific markers. smFISH revealed selective and extensive co-localization of *Cadps2* with *Calca* or with *Fst* transcripts within the same subpopulation of cells in the MC layer **(Figure 3G and G’, Figure 3-figure supplement 1N)**. Furthermore, *Cadps2, Calca* and *Fst* expression was consistently observed in neurons expressing GFP **(Figure 3G and H, Figure 3-figure supplement 1K-M)**. Importantly, two-color smFISH revealed that type-specific M1, M2 and M3 markers were expressed in largely non-overlapping populations of MCs: M1-specific *Kcng1* and *Lhx5* transcripts did not co-localize with M2-specific *Vgll2* and *Piezo2* transcripts (**Figure 3-figure supplement 1O and P**); M1-specific *Kcng1* transcripts did not co-localize with M3-specific *Cadps2* and *Calca* transcripts **(Figure 3H and H’, Figure 3-figure supplement 1Q)**; M2-specific *Vgll2* transcripts did not co-localize with M3-specific *Cadps2* transcripts **(Figure 3I and I’)**. Together, the selective co-localization of type-specific genes in non-overlapping populations in the mitral cell layer validates these three types as accurate and meaningful groupings of MCs, and their co-localization with GFP validates their identity as projection neurons.

DE analysis for TC type-specific genes identified the transcription factor BarH-like homeobox 2 (*Barhl2)*, the gamma-sarcoglycan *Sgcg*, the vitamin D receptor (*Vdr)*, and the olfactory receptors *Olfr110* and *Olfr111* as putative T1 markers. Two-color smFISH revealed extensive co-localization of *Barhl2* and *Sgcg* or *Olfr110/Olfr111* transcripts within the same subpopulation of cells in the external plexiform layer, indicative of middle tufted cells **(Figure 3J and J’, Figure 3-figure supplement 2B)**. Furthermore, *Barhl2* and *Sgcg* expression was observed in neurons expressing GFP **(Figure 3J)**.

The coagulation factor C homolog (*Coch)* and the Wnt family member 5b (*Wnt5b)* were identified as putative ET1 markers. smFISH confirmed the expression of *Coch* and *Wnt5b* in a subpopulation of cells in the external plexiform and glomerular layers **(Figure 3K, figure supplement Figure 2D)**. Moreover, *Coch* expression was observed in neurons expressing GFP **(Figure 3K)**. The LIM homeobox 1 (*Lhx1)* and the early B-cell factor 3 (*Ebf3)* were identified as putative ET2 markers. Two-color smFISH revealed *Lhx1* and *Ebf3* co-expression in a sparse subpopulation of cells at the boundary between the external plexiform and glomerular layers **(Figure 3-figure supplement 2F)**, indicative of external TCs. Finally, we identified the lymphocyte antigen 6 family member 6GE (*Ly6g6e)* and the transcription factor Forkhead box O1 (*Foxo1)* as putative ET4 markers. smFISH revealed selective expression of *Ly6g6e* and *Foxo1* in a subpopulation of cells in the glomerular layer **(Figure 3-figure supplement 2H)**, and co-localization of *Ly6g6e* with GFP validates their identity as external TCs **(Figure 3L)**. Importantly, two-color smFISH revealed that type-specific tufted cell markers were expressed in largely non-overlapping populations of cells: ET1-specific *Coch* transcripts did not co-localize with T1-specific *Barhl2* transcript or with ET4-specific *Ly6g6e* transcripts **(Figure 3K and K’, 3L and L’)**. Furthermore, ET4-specific *Foxo1* and *Ly6g6e* transcripts did not co-localize with T1-specific *Barhl2* transcript **(Figure 3-figure supplement 2I)**, with ET2-specific *Ebf3* transcript **(Figure 3-figure supplement 2J)** or with ET1-specific *Wnt5b* transcript **(Figure 3-figure supplement 2K)**. Overall, the selective co-localization of type-specific genes, their location within the olfactory bulb, their non-overlapping nature, and their co-localization with GFP validates these five types of middle and external TCs as accurate and meaningful classifications.

### Inferring gene regulatory networks for projection neurons

The differential gene expression patterns revealed by transcriptome analysis are determined by the concerted action of transcription factors (TFs). We therefore set out to characterize cell types by their TF activity, and we used independent information about TF binding sites to group genes by TF interactions. These gene regulatory networks are biologically meaningful and more robust against technical artifacts than the expression of individual genes, providing a complementary set of axes along which to cluster MCs and TCs. Ultimately, gene regulatory network analysis can yield more detail for classifying cell types and for understanding the molecular mechanisms that underlie their transcriptional differences.

To infer the regulatory networks of each type of projection neuron, we used the Single-Cell Regulatory Network Inference and Clustering pipeline (SCENIC, (Aibar et al., 2017)). SCENIC is a three-step computational protocol based around regulons. A regulon is a TF and its (predicted) target genes **(Figure 4A)**. In brief, SCENIC consists of co-expression analysis, followed by TF binding motif enrichment analysis, and finally evaluation of a regulon’s activity. The results are a list of regulons and a matrix of all the single cells with their regulon activity scores (RAS, essentially an Area-Under-the-Curve metric, see Methods for details).

**Figure 4:**
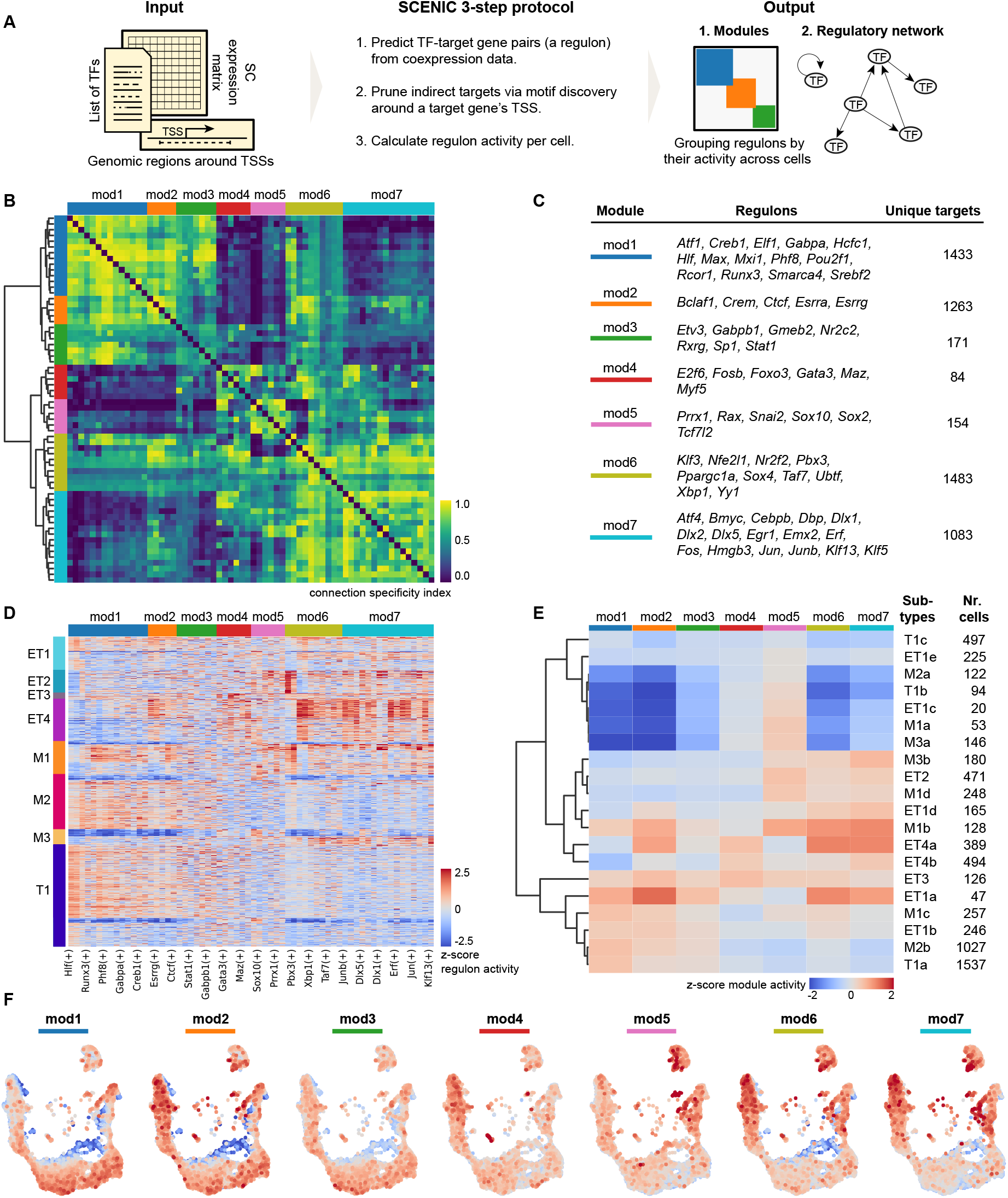
Mitral and tufted cell-specific regulons combine into modules. **(A)** Schematic representation of the network analysis pipeline, including the required input, the SCENIC protocol, and the output in the form of regulon modules and a regulatory network **(Figure 5)**. **(B)** Hierarchical clustering of regulons using the Connection Specificity Index (CSI) as a distance measure results in 7 modules (Ward linkage). The CSI of two regulons is based on Pearson correlation coefficients (PCC): for each PCC between regulon A and B, the CSI is the fraction of regulons that have a PCC with A and with B lower than PCC (AB). Prominent cross-module interactions are observed for mod1-2-3 and mod6-7. Moreover, module 6 showed interactions with all other modules. **(C)** Table listing the modules, their regulons, and the number of unique target genes in each module. **(D)** Projection neuron cell types defined by transcriptome analysis **(Figure 2)** and subtypes (rows) defined by regulon activity (columns). Rows were ordered by cell type and within a cell type by hierarchical clustering (Euclidean distance, Ward linkage). Columns clustered as in panel **B**. **(E)** Module activity per cell subtype. Module activity is calculated as the average activity of its regulons for a given cell subtype. Each cell subtype may be defined by a combination of active and inactive modules. For example, T1a (bottom row) is defined by high activity in modules 1, 2, and 3. **(F)** Module activity mapped on the transcriptome UMAP space. Color range as in panel **E**.

### Clustering on regulon activity corroborates molecular groupings of mitral and tufted cell types and allows further subdivision

We applied SCENIC to MCs and TCs (6472 cells), computing 64 regulons with a range of 8 to 724 target genes (median = 41). These regulons greatly reduce the dimensionality of the data from >30,000 genes to 64 regulons, defining a new biologically meaningful space in which to analyze relationships between cells. Using the regulon activity matrix, we performed a Leiden clustering (Traag et al., 2019; Wolf et al., 2018) in UMAP space on the putative projection neurons and confirmed that doing so recapitulated the cell types we defined based on transcriptome analysis **(Figure 4-figure supplement 1)**. We observed that the two approaches produced similar clusters, with only minor differences in drawing the boundaries between clusters **(Figure 4-figure supplement 1)**. The strong overlap between these classification methods validates the cell types we characterize as meaningful divisions within the data. In addition, constraining genome-wide transcriptome data by TF-target gene interactions shows that the transcriptome-defined MC and TC types are heterogenous clusters in terms of regulon activity **(Figure 4D)**. This suggests the existence of subtypes within mitral and tufted cell types. To investigate these further, we defined more fine-grained groupings within cell types. We used hierarchical clustering analysis to further subdivide cell types, finding 5 subtypes of M1, 2 subtypes each of M2 and M3, 5 subtypes each of T1 and ET1, and 3 subtypes of ET4. We confirmed that these subtypes were well-recovered with a Leiden clustering on regulon activity **(Figure 4-figure supplement 2)**.

### Combinations of regulon modules characterize mitral and tufted cell subtypes

TFs activity is thought to be organized into coordinated network modules that determine cellular phenotypes (Alexander et al., 2009; Irons and Monk, 2007; Suo et al., 2018). To characterize how TFs are organized into such modules in OB projection neurons, we searched for combinatorial patterns of regulon activity. We used the Connection Specificity Index (CSI) to this end, which is an association index known to be suited for the detection of modules (Fuxman Bass et al., 2013; Suo et al., 2018). By computing the CSI on the basis of pairwise comparisons of regulon activity patterns across cells, we found that the 68 regulons grouped into 7 modules (mod1-7) **(Figure 4B and C)**. We confirmed these modules by looking at the activity of individual regulons in each cell and visually verifying that individual regulons act together as the identified modules **(Figure 4D)**.

Interestingly, regulon and module activity were not uniform within cell types. Rather, regulon activity suggested distinct subtypes of each MC or TC type **(Figure 4D)**, corroborating that using biologically relevant information to reduce dimensionality facilitates more fine-grained classification. To further investigate these subtypes, we used the modules to describe how the combined regulatory logic of distinct TFs contributes to the diversity of MC and TC subtypes. We asked if combinations of modules could uniquely describe the subtypes. To do so, we calculated the average module activity score per cell subtype. Next, we performed a hierarchical clustering (correlation distance, complete linkage) on the subtypes **(Figure 4E)** and mapped average module activity on the UMAP space defined by transcriptome analysis **(Figure 4F)**. We found that, when grouped by module activity, subtypes do not group by type; rather, subtypes of different cell types share similar module activity **(Figure 4E)**, providing another complementary axis for grouping cells. Interestingly, module activity also forms gradients along the UMAP plot, representing gradual transitions between subtypes **(Figure 4F)**. Therefore, while subtypes were identified most easily using modules of regulons, this subtype structure is also apparent in the full transcriptome space, further validating the divisions between subtypes as accurate and meaningful. Moreover, it makes explicit that a continuous activity gradient of TFs is transformed in a non-linear manner into distinct transcriptome differences between mitral and tufted cell types.

### Regulon-based transcription factor networks reveal overlapping features of cell type identity

While TFs regulate large numbers of target genes, central to cell identity are the interactions between TFs: TFs can regulate their own expression as well as the expression of other TFs, generating a TF network thought to be a core determinant of cell type identity (Arendt et al., 2016; Becskei et al., 2001; Thieffry, 2007). We thus looked at TF-TF interactions to visualize the TF network topology that defines MCs and TCs. We specifically asked whether MC and TC classes share common TF network features, or whether, as suggested by the analysis of genome-wide transcriptome and regulon analysis **(Figures 2-4)**, MC and TC subtypes are defined by specific yet overlapping TF network features.

As a regulon is defined by a TF and a set of target genes, we constructed a (directed) network of TFs by taking from each regulon’s target genes only the TFs that have regulons themselves **(Figure 5A)**. Overall, we found one large set of interconnected TFs, three small components of five TFs or fewer, and twelve isolated TFs. 38 of the 64 TFs (59%) show possible self-activating regulation, and several others form mutually-activating pairs (e.g. *Mxi1* and *Phf8*). The network is dominated by three hub genes, two of which may self-activate: *Pbx3* (activates 19 TFs), *Bmyc* (activates 10 TFs) and *Bclaf1* (activates 7 TFs). Their central position in this network suggests a role as hub regulators of MC and TC identity.

**Figure 5:**
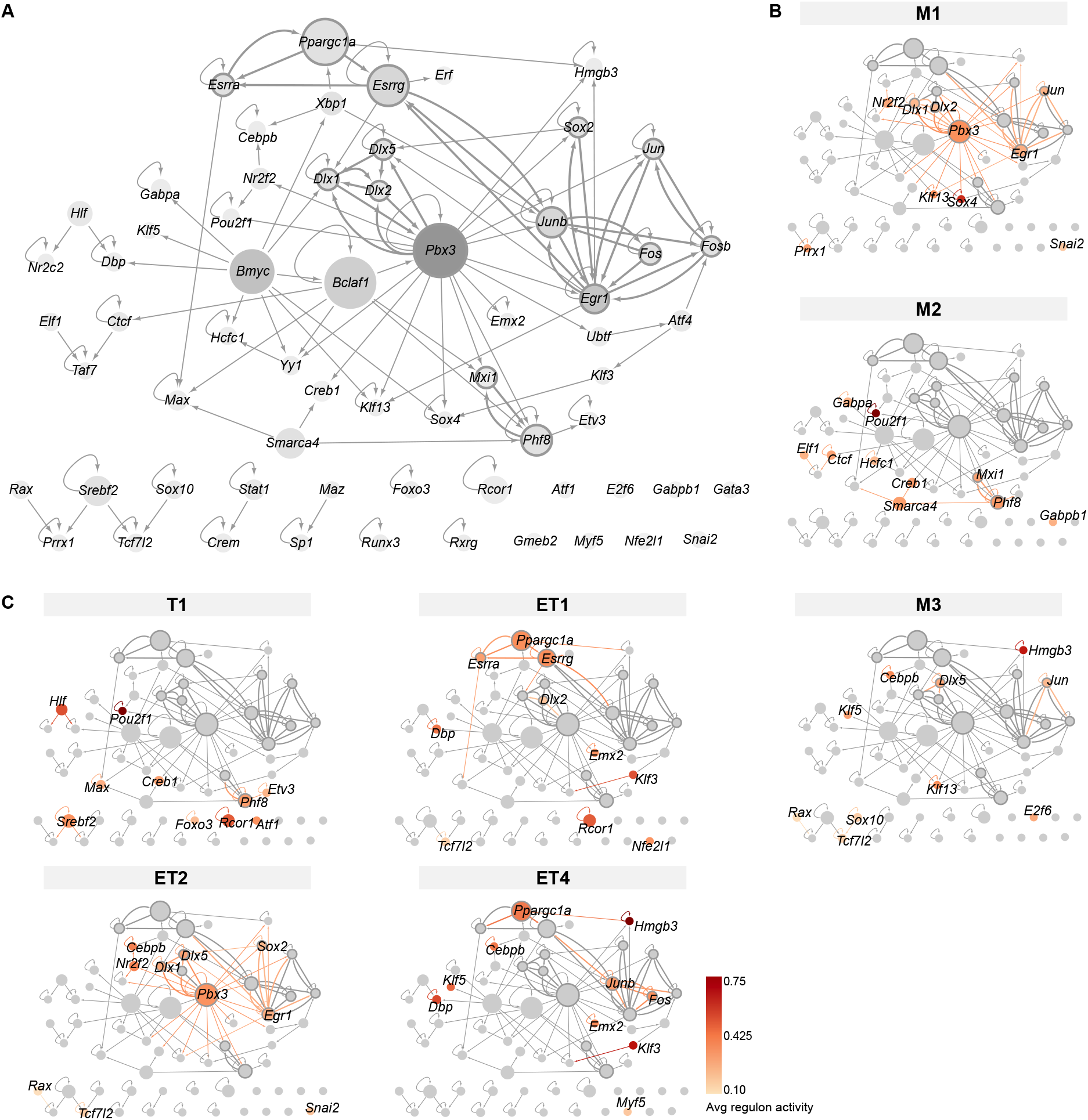
Transcription factor network derived from regulons. **(A)** Overview of mitral and tufted cell transcription factor (TF) networks, with node size scaled by the number of target genes and nodes colored with different shades of gray based on outdegree (number of outgoing edges). Thick borders and edges denote cycles of 2 or 3 regulons. The three main hubs are: *Pbx3* (outdegree 19, target genes 688), *Bmyc* (outdegree 10, target genes 578) and *Bclaf* (outdegree 7, target genes 724). **(B)** Mitral cell types (M1, M2, M3) with standardized regulon activity for the top 10 most specific regulons mapped onto the corresponding TF nodes (compare **Figure 4D, E**). **(C)** Tufted cell types (T1, ET1, ET2, ET4) with standardized regulon activity for the top 10 most specific regulons. We omitted ET3 as it only had a few cells.

As anticipated from our module analysis, we find features that are shared across certain types of MCs and TCs rather than MC- or TC-specific network features **(Figure 5B and C)**. For instance, we find that M1 and ET2 are characterized by the hub *Pbx3* and *Dlx1*, which are both self-activating and together form a positive feedback loop. Yet both cell types are distinguishable by additional cycles: M1 has *Dlx2* “in the loop” with *Pbx3* and *Dlx1*, whilst ET2 has a 5-cycle involving *Pbx3, Dlx1, Dlx5, Sox2* and *Egr1* (i.e. a superset of the shared cycle with M1). For M2 and T1, we observe that they are mostly characterized by peripheral nodes of the network, that in most cases do not regulate each other, even if many are grouped together in module 1 **(Figure 4C)**. And ET1 and ET4 are characterized by the target-gene-rich 3-cycle of *Ppargc1a* and the estrogen receptors *Esrra, Esrrg* (for ET4 the estrogen receptors are in the top 20 of specific regulons), with ET4 also featuring the cycle between *Junb* and *Fos*. Interestingly, these two cycles are connected, for instance via *Esrrg* and *Junb*.

Taken together, the analysis of TF regulatory networks suggests that individual MC and TC types share key TF network features, which might point towards common physiology or connectivity features. The differentially active TF network hubs and loops provide starting points for future investigation of the functional differences between the MC and TC types described here. Thus, the modules and the network serve as complementary approaches for studying cell type identity, with modules suited to classifying cells into types and subtypes and network analysis suited to investigating their functional differences.

### Simulating single nucleus gene expression from bulk RNA deep sequencing

TCs preferentially target anterior olfactory regions, including the Anterior Olfactory Nucleus (AON), while MCs target anterior and posterior olfactory areas (Imamura et al., 2020). Therefore, we asked whether the genetic diversity between MC and TC types could provide information about their projection targets. To investigate this question, we again injected rAAV-retro-CAG-H2B-GFP into olfactory cortex, albeit now *either* into the AON *or* the posterior Piriform Cortex (pPCx) **(Figure 1A)**. For each injection site and in three replicates, we then enriched GFP-expressing OB nuclei, and prepared RNA for bulk RNA deep sequencing **(Figure 1A)**.

Bulk RNA sequencing represents molecular information from a variety of different cell types. Given that a substantial fraction of isolated nuclei in our experiments was comprised of granule and periglomerular cells, in addition to projection neurons, we devised a novel computational approach to recapitulate the constituent cell types from bulk RNA sequencing data by simulating single nucleus expression profiles. Previous methods simulated the transcriptome of a single cell based on the overall distribution of gene expression levels in the bulk RNA sequencing data, producing many nuclei that were similar to each other and to the original bulk expression profile (Konstantinides et al., 2018). This worked well for clean bulk RNA-seq datasets with only one cell type, but for our mixed datasets, the simulated nuclei resembled an unrealistic average of the constituent cell types. Therefore, to capture the diversity contained within our bulk RNA-seq datasets, we used regulons as the unit of analysis to create simulated nuclei with more biologically realistic transcriptomes (**Figure 6A and B**).

**Figure 6:**
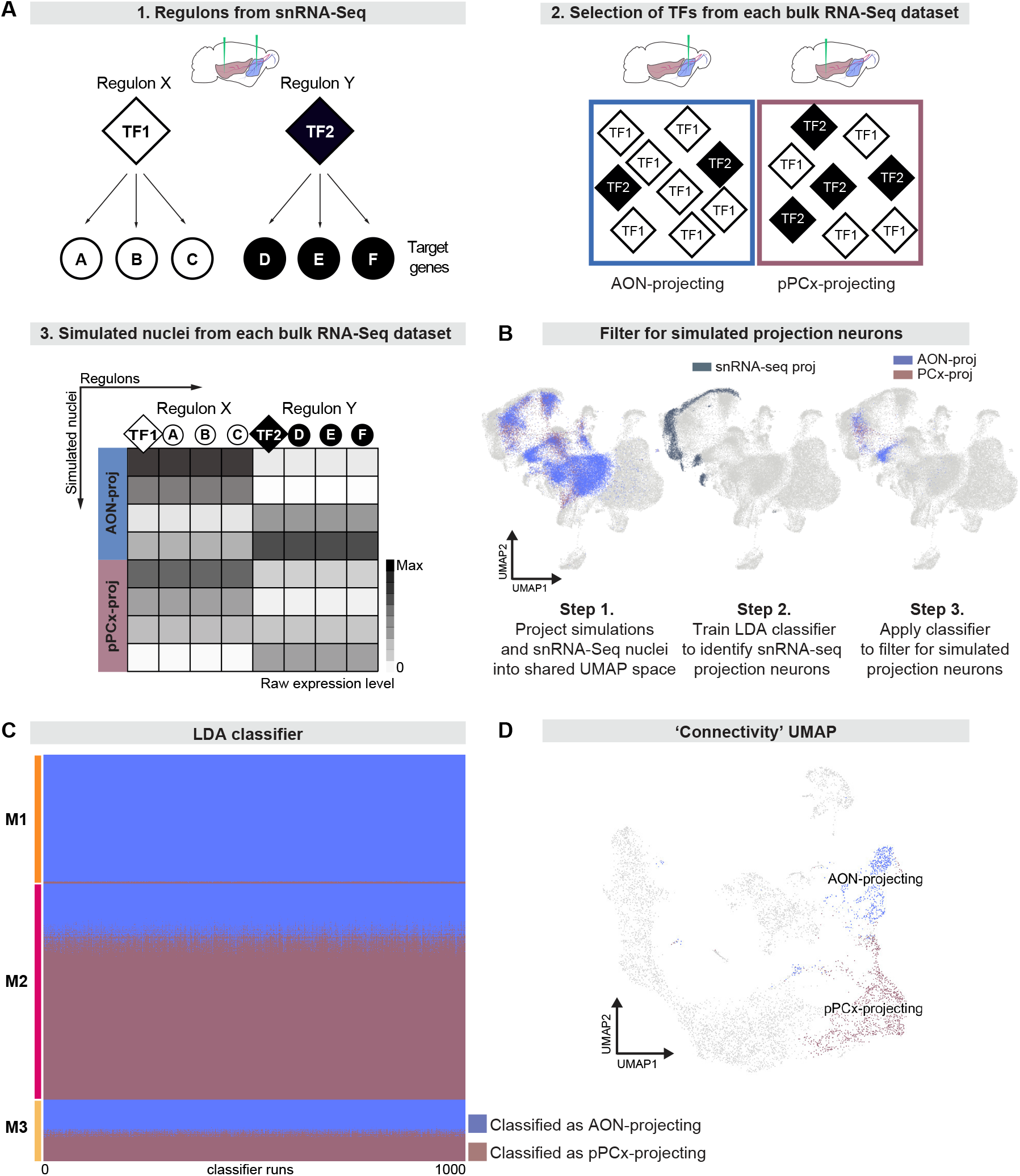
Simulations from bulk RNA deep sequencing data suggest mitral types have distinct projection targets. **(A)** Schematic representation of strategy to integrate bulk RNA-seq and single nucleus RNA-seq data. **(A1)** Simulations use regulons inferred from snRNA-seq data. A regulon consists of a transcription factor and all target genes that are activated by that transcription factor. **(A2)** When simulating nuclei from bulk RNA-seq data, the probability of selecting a given regulon is determined by the abundance of its transcription factor in the bulk RNA-seq dataset. **(A3)** Nuclei are simulated from each bulk dataset through random sampling of regulons with replacement. This method maintains broad differences between datasets while accounting for heterogeneity within each dataset. **(B)** Simulated nuclei and snRNA-seq nuclei projected into a shared UMAP representation. Step 1: Blue indicates all simulated nuclei from AON-projecting bulk RNA-seq and purple indicates all simulated nuclei from pPCx-projecting bulk RNA-seq. Step 2: Darker color indicates snRNA-seq projection neurons. Step 3: Blue indicates AON-projecting simulated projection neurons and purple indicates pPCx-projecting simulated projection neurons. **(C)** Linear Discriminant Analysis (LDA) classifiers were trained on simulated projection neurons, then used to predict the projection target of snRNA-seq mitral cells to investigate projection targets of snRNA-seq derived types. Each row represents one mitral cell. Each column represents one of 1000 LDA classifiers. Blue indicates that the mitral cell was classified as AON-projecting, and purple indicates that the mitral cell was classified as pPCx-projecting. Within each mitral cell type, cells are sorted vertically by their predicted projection target. **(D)** UMAP representation color-coded by predicted projection target. Cells in blue were predicted to be AON-projecting by all 1000 classifiers. Cells in purple were predicted to be pPCx-projecting by all 1000 classifiers.

We first compared simulated and snRNA-seq nuclei by using principal component analysis to project both into a shared low-dimensional space. We used these principal components as input to a UMAP projection to visually inspect relationships between simulated and snRNA-seq nuclei (**Figure 6B**). Consistent with histology and snRNA-seq analyses (**Figures 1 and 2**), simulations from both AON-injected and pPCx-injected bulk RNA-seq datasets contained cells types other than projection neurons, and this contamination was more pronounced in simulations from AON-injected bulk RNA-seq datasets. The dispersion of simulated nuclei throughout this space indicated that simulations successfully recapitulated the diversity of cell types in the bulk RNA-seq datasets, as each cell class from the combined snRNA-seq data had some simulated nuclei in its vicinity (**Figure 6B**). To account for this contamination and filter for only putative simulated projection neurons, we trained linear discriminant analysis (LDA) classifiers to predict whether snRNA-seq nuclei were projection neurons based on the 30 top principal components that defined the shared low-dimensional space. These classifiers accurately and consistently classified projection neurons, with a mean Jaccard index (a measure of similarity between predicted and true labels) of 97.3% and a standard deviation of 0.17% over 1000 classifiers. We then applied this classifier to the simulated nuclei, designating those simulations predicted to be projection neurons by all 1000 classifications as putative simulated projection neurons (**Figure 6B**).

To directly compare these simulated projection neurons to MCs characterized through snRNA-seq, we next used principal component analysis to define a shared low-dimensional space for MCs and simulated projection neurons only. To investigate potential differences in projection target between MC types, we trained 1000 LDA classifiers to predict the projection target of simulated projection neurons based on the 30 top principal components that defined the shared low-dimensional space (mean Jaccard index: 85.1%; standard deviation: 1.1%). We then used these classifiers to predict the projection targets of snRNA-seq MCs. Interestingly, we consistently found different predicted targets for the molecularly-defined MC types. 98.4% of M1 MCs were classified as AON-projecting by at least 90% of classifiers, suggesting that M1 cells preferentially project to anterior targets (**Figure 6C**). In contrast, 71.4% of M2 MCs were consistently classified as pPCx-projecting, suggesting that M2 cells preferentially project to posterior targets (**Figure 6C**). M3 MCs segregated into distinct populations: 48.2% of M3 MCs were consistently classified as AON-projecting, while 42.9% were consistently classified as pPCx-projecting (**Figure 6C**). These findings suggest that the molecular subcategorization of MCs may delineate differences in connectivity. For M1 and M2, these labels were defined by gene expression and regulon activity, and projection target specificity. However, in the case of M3, our results suggest that the molecular category contains neurons with distinct projection patterns (**Figure 6D)**. These results suggest that neuronal connectivity provides an independent axis along which to investigate cell type identity.

## Discussion

Morphological differences between OB mitral and tufted cells have been described since the time of Cajal (The Croonian lecture, 1894). Electrophysiological and functional imaging experiments *in vivo* and *in vitro*, developmental studies as well as anatomical reconstructions from light and electron microscopy studies have further highlighted the heterogeneity of OB projection neurons (Christie et al., 2001; Ezeh et al., 1993; Fukunaga et al., 2012; Geramita et al., 2016; Haberly and Price, 1977; Kawasawa et al., 2016; Mori et al., 1983; Phillips et al., 2012). We here provide the first detailed molecular profiling of projection neurons of the mouse olfactory bulb and delineate the types and subtypes of mitral and tufted cells together with their key gene regulatory networks.

We have performed single nucleus and bulk RNA deep sequencing to characterize the molecular diversity of mouse OB projection neurons. We identified, based on transcriptome and RNA *in situ* analysis, three distinct types of MCs and five distinct types of TCs. We then used comprehensive gene regulatory network analysis to reveal candidate gene regulatory mechanisms that underlie cell type diversity. Finally, we describe a novel computational approach for integrating single nucleus and bulk RNA sequencing data, and we use this approach to propose that different MC types selectively project to anterior versus posterior regions of olfactory cortex. Our analyses identified potential molecular determinants of cell type-specific functional properties and projection target connectivity and provide a comprehensive resource for investigating odor processing and olfactory circuit function and evolution.

### The molecular diversity of olfactory bulb projection neurons

Given that the vast majority of OB neurons are interneurons, notably granule cells and juxtaglomerular neurons, we devised a retrograde viral targeting strategy to substantially enrich for OB projection neurons. This allowed us to analyze the transcriptomes of over 7500 putative projection neurons that could in turn be grouped into 8 molecularly distinct mitral and tufted cell types. We validated neuronal cell identity using smFISH with multiple type-specific marker genes, and we determined the localization of identified neuronal types within the mitral cell, external plexiform and glomerular layers of the olfactory bulb. Finally, we used retrograde viral tracing to confirm that the molecularly distinct neuronal types we describe indeed project to the olfactory cortex. Based on our analysis we define three molecularly distinct types of MCs and five distinct types of TCs.

The neuronal cell types we have characterized here likely represent the major categories of OB projection neurons. More extensive sampling might reveal additional rare cell types, and more fine-grained clustering could further subdivide subtypes of neurons. However, our samples contained 7,504 putative projection neurons compared to current estimates of 10,000 – 30,000 projection neurons overall per OB (Nagayama et al., 2014; Richard et al., 2010). Furthermore, we sorted nuclei rather than whole cells, which is thought to more accurately reflect relative cell type abundance (Habib et al., 2017; Lake et al., 2017). Altogether we are therefore confident that our analysis captures the key biologically relevant types of projection neurons. Independent from the number of molecularly distinct neuronal cell types, the gene expression profiles we have described here provide critical new tools for refining projection neuron cell type identities by aligning a cell’s molecular features with its functional properties. Previous experiments have highlighted the heterogeneity of odor responses of MCs and TCs (Balu et al., 2004; Bathellier et al., 2008; Carey and Wachowiak, 2011; Desmaisons et al., 1999; Friedman and Strowbridge, 2000; Schaefer et al., 2006). We propose that this functional diversity can be explained, at least in part, by the molecular diversity of OB projection neurons, a model that can now be tested experimentally.

### Specificity of mitral and tufted cell projections

A critical feature of neuronal cell type identity is their projection target specificity. Earlier studies have shown that TCs project to anterior regions of the olfactory cortex only, while MCs project to both anterior and posterior olfactory cortex (Imamura et al., 2020; Nagayama et al., 2010; Scott et al., 1980). Furthermore, and in contrast to the organization of neuronal projections to sensory cortexes for vision, hearing, or touch, projections from the OB to the piriform cortex lack apparent topographical organization (Ghosh et al., 2011; Miyamichi et al., 2011; Sosulski et al., 2011).

We have analyzed, using bulk RNA deep sequencing, gene expression profiles of cells projecting to anterior versus posterior olfactory cortex. Using simulations based on gene regulatory network analysis, we have then mapped the bulk RNA sequencing data onto MC types defined by single nucleus RNA sequencing. Interestingly, our analysis suggests that cells of the M1 MC type preferentially target the anterior olfactory cortex, while M2 cells preferentially target the posterior olfactory cortex. Furthermore, M3 cells, while clustering as a single cell type based on their transcriptomes, project to either anterior or posterior cortical targets. These results are consistent with the model that gene expression and connectivity provide complementary axes for cell type classification (Kim et al., 2020). Importantly, our results provide critical new molecular markers for investigating the anatomical organization of connectivity from the OB to cortex.

### Molecular mechanisms underlying olfactory bulb projection neuron diversity

Gene regulatory network analysis can reveal the transcriptional programs that determine the functional properties of neuronal subtypes. Here, we describe cell type-specific modules of gene regulation, defined by the interactions of transcription factors and their target genes. One intriguing result of this analysis is that cell subtypes do not fall into clearly delineated MC and TC classes. For example, module activity of M1d and M3b MC subtypes is more similar to those of ET2 TCs than other M1 and M2 MC subtypes. We obtained similar results from analyzing the TF-TF network that is thought to be closely linked to maintenance of cell identity. This demonstrated that while some MC subtypes indeed share key TFs with each other, regulon activity and network features in MC subtypes and TC subtypes are highly overlapping. For example, the most prominent hub, *Pbx3*, is strongly active in M1 and ET2 but not any of the other MC or TC subtypes. Generally, projection neurons are characterized by a variety of hubs and cycles in the TF-TF network that are used by both MC and TC subtypes in a combinatorial manner. The prominent hub and cycle-related genes we have identified here, may act as candidate master regulators of neuronal function, which can be targeted for experimental validation. Together, these results suggest that subtypes of MCs and TCs may share important functional properties, possibly blurring at the transcriptional level the traditional division into tufted and mitral cells as the two major classes of OB projection neurons. Moreover, the gradients of module activity that we observed over the MC and TC subtypes theoretically provide a mechanism for generating multiple distinct cellular phenotypes, similar to how morphogenetic gradients allow for spatial patterning and cell differentiation (Wolpert, 1969). Through non-linear regulatory interactions gradual differences at the transcriptomic level can be translated into selective expression of functional genes.

For example, we found that the *Kcng1* gene was selectively expressed in M1 MCs. The *Kcng1* gene encodes for a voltage-gated potassium channel, which forms heterotetrameric channels with the ubiquitously expressed delayed rectifying Kv2.1 potassium channel (indeed also expressed in the M1 cluster) and modifies the kinetics of channel activation and deactivation (Kramer et al., 1998). Other voltage-gated potassium channels exhibiting prominent differential expression levels in MC and TC subtypes include *Kcnd3, Kcng1, Kcnh5, Kcnq3, Kcnj2* and 6, and *Hcn1* (for details see accompanying website link in Materials and Methods). These channels represent intriguing candidates for controlling the differential excitability of different MC and TC types.

We also found that a large number of cell adhesion and axon guidance genes known to control the formation and maintenance of neuronal connectivity were differentially expressed in OB projection neuron types. Examples include members of the cadherin superfamily of cell adhesion glycoproteins (*Cdh6, 7, 8c, 9, 13*, and *20*), and components of the Semaphorin/Neuropilin/Plexin complexes, including *Nrp2, Plexna3, Sema3e*, and *Sema5b*. Semaphorin/Neuropilin/Plexin complexes are known to play critical roles in the development and maintenance of neuronal connections, including in OB MCs (Inokuchi et al., 2017; Saha et al., 2007). Heterogeneity in these cell adhesion and guidance genes might inform subdivisions in projection neurons across the OB, in particular along the dorsomedial-ventrolateral axis.

Our data set provides an important resource for studying the evolution of olfactory neural circuits across species. Adaptation to distinct olfactory environments, and the critical role of olfaction in survival and reproduction has shaped the evolution of the repertoire of odorant receptors and olfactory sensory neurons (Bargmann, 2006; Niimura, 2012). However, little is known about how evolving sensory inputs from the nose are accommodated at the level of the OB and its connections to the olfactory cortex. A detailed molecular description of mouse OB projection neurons provides a first step towards understanding the evolution of olfactory sensory processing across species.

## Materials and methods

### Experimental model and subject details

Male and female C57Bl/6 mice (6- to 8-week-old) were used in this study and obtained by in-house breeding. All animal protocols were approved by the Ethics Committee of the board of the Francis Crick Institute and the United Kingdom Home Office under the Animals (Scientific Procedures) Act 1986, as well as Brown University’s Institutional Animal Care and Use Committee followed by the guidelines provided by the National Institutes of Health.

### Stereotaxic injections and histology

Mice were anaesthetized using isoflurane and prepared for aseptic surgery in a stereotaxic frame (David Kopf Instruments). A retrogradely transported Adeno Associated Virus (rAAV-retro-CAG-H2B-GFP, (Tervo et al., 2016)) was injected stereotaxically into multiple sites of piriform cortex (PCx) and anterior olfactory nucleus (AON). The following coordinates, based on the Paxinos and Franklin Mouse Brain Atlas, were used: Coordinates (AP / ML / DV) in mm for PCx injections: (1) −0.63 / −4.05 / −4.10, (2) −0.8 / −4.00 / −4.10. For AON injections: (1) 2.8 / 1.25 / 2.26 and 2.6, (2) 2.68 / 1.25 / 2.3 and 2.75, (3) 2.34 / 0.7 / 3.5. Using a micromanipulator, a pulled glass micropipette was slowly lowered into the brain and left in place for 30 seconds before the virus was dispensed from the micropipette using a Nanoject injector (Drummond Scientific) at a rate of 46 nl/min (0.3 μl for PCx and 0.2 μl for AON per injection site). The micropipette was left in place for an additional 5 min before being slowly withdrawn to minimize diffusion along the injection tract. Craniotomies were covered with silicone sealant (WPI) and the skin was sutured. Mice were provided with 5 mg/kg Carprofen in their drinking water for 2 days following surgery.

Histology was used to validate viral targeting of olfactory bulb projection neurons. Mice were deeply anaesthetized with 2.5% of 250mg/kg Avertin and transcardially perfused with 10 ml of ice-cold phosphate-buffered saline (PBS) followed by 10 ml of 4% paraformaldehyde (PFA). Brains were dissected and post-fixed for 5 h in 4 % PFA at 4 °C. Coronal sections (100 µm thick) were prepared using a vibrating-blade Leica VT100S Vibratome. Sections were rinsed in PBS and incubated in PBS / 0.1% Triton X-100 and Neurotrace counterstain (1:1000, ThermoFisher) at 4 °C overnight, then mounted on SuperFrost Premium microscope slides (Fisher, cat# 12-544-7) in Fluorescent Vectashield Mounting Medium (Vector). Images were acquired at 20X using a Nikon A1R-HD confocal microscope.

### Single nuclei isolation, FANS and RNA extraction

To isolate GFP-labeled nuclei, 9 individual replicates were used (for bulk RNA deep sequencing: 3 replicates of AON-injected mice and 3 replicates of PCx-injected mice; for single nuclei RNA sequencing: 3 replicates of AON and PCx-injected mice). Mice were deeply anaesthetized with an overdose of ketamine/xylazine and transcardially perfused with ice-cold PBS. Both hemispheres of the olfactory bulb were dissected, and the hemisphere ipsilateral to the injection site was carefully separated from the contralateral hemisphere. Both hemispheres were minced separately and placed into two different tubes. To dissociate single nuclei, Nuclei PURE Prep was used according to the manufacturer instructions (Sigma, cat# NUC201-1KT) with some modifications. The minced tissue was gently homogenized in 2.75 ml Nuclei PURE Lysis Buffer and 27.5 μl 10% Triton X-100 using an ice-cold dounce and pestle, and filtered two times through a 40 μm cell strainer on ice. After centrifuging at 500 rpm for 5 min at 4 °C, the supernatant was aspirated and gently resuspended in 500 μl of cold buffer (1x of cold Hanks’ Balanced Salt Solution HBSS, 1% nuclease-free BSA, 22.5 μl of RNasin Plus (Promega N2611) and 1/2000 DRAQ5).

Fluorescence-activated nuclei sorting of single nuclei was performed using a BD FACSAria™III Cell Sorter with a 70 μm nozzle at a sheath pressure of 70 psi. Precision mode (yield mask set to 16, purity mask set to 16 and phase mask set to 0) was used for stringent sorting. For single nucleus RNA sequencing, GFP+ nuclei were sorted into a 1.5 ml centrifuge tube. For bulk RNA deep sequencing, GFP+ nuclei were sorted into 100 μl Trizol and 1.43 μl of RNA carrier, and total RNA was extracted using the Arcturus PicoPure RNA Isolation Kit (ThermoFisher, cat# KIT0204).

### Single nucleus RNA sequencing

Libraries were prepared using the Next Single Cell / Low Input RNA Library Prep Kit (New England Biolabs). The quality and quantity of the final libraries were assessed with the TapeStation D5000 Assay (Agilent Technologies) before sequencing with an Illumina HiSeq 4000 platform using the 10X kit version Chromium Single Cell 3’ v3. RNA concentrations were measured as: 14.4, 23.3, 7.9 ng/μl (n = 3 animals).

### Bulk RNA deep sequencing

Libraries were prepared using the Next Single Cell / Low Input RNA Library Prep Kit (New England Biolabs). The quality and quantity of the final libraries was assessed with the TapeStation D1000 Assay (Agilent Technologies) before sequencing with an Illumina HiSeq 4000 platform. RNA concentrations were measured as: AON injections (n = 3 animals), 1.495, 1.682, 1.881 ng/μl and RNA integrity numbers (RIN) 8.3, 8.7, 9.0; PCx injections (n = 3 animals), 0.257, 0.165, 0.133 ng/μl; RIN = 8.0, 10.0, 7.8 for each replicate, respectively.

### Single nucleus RNA sequencing analysis

Raw sequencing datasets were processed using the Cell Ranger pipeline (10x Genomics). Count tables were loaded into R (version 3.6, https://www.r-project.org) and further processed using the Seurat 3 R-package (Butler et al., 2018).

For each of the three replicates, we removed all nuclei with fewer than 500 distinct genes detected or with more than 5% of unique molecular identifiers stemming from mitochondrial genes. After quality control, we merged the replicates and retained a total of 31,703 nuclei (median of 2,300 genes per nucleus; for each replicate median genes per nucleus: R1=2,266; R2=2419; R3=2,322). Principal component analysis (PCA) was then performed on significantly variable genes and the first 30 principal components were selected as input for clustering and UMAP, based on manual inspection of a principal component variance plot (PC elbow plot). Clustering was performed using the default method (Louvain) from the Seurat package, with the resolution parameter of the FindClusters function set to 0.3.

Subclustering of projection neurons was carried out by selecting clusters M1, M2/M3, T1, ET1 and ET2 from the initial single-nuclei analysis based on the combinatorial expression patterns of glutamatergic and previously characterized mitral/tufted cell markers (*Tbx21, Pcdh21, Thy1, Vglut1, Vglut2 and Vglut3)*. Subclustered nuclei were subjected to a new clustering with the Seurat resolution parameter of the FindClusters function set to 0.3.

Differential gene expression analysis on single-nuclei data was performed using the glmGamPoi R-package (Ahlmann-Eltze and Huber, 2020). Gene set enrichment analysis (GSEA) on the resulting log-fold changes was performed as described in (Subramanian et al., 2005).

### Network inference

Gene regulatory networks were inferred using the pySCENIC pipeline (Single-Cell rEgulatory Network InferenCe, (Aibar et al., 2017)) and visualized using Jupyter notebooks and Cytoscape (Shannon et al., 2003). pySCENIC is a three-step approach: (1) predict TF-target gene pairs using Arboreto; (2) filter TF-target gene associations for false positives using TF binding site enrichment in a window of 5kb around a target’s Transcription Start Site (TSS) and group TFs with their target genes into so-called regulons; (3) calculate the activity of regulons in each cell in terms of the Area Under the recovery Curve (AUC). Step 1 depends on a stochastic search algorithm and is therefore performed n = 100 times. Only TFs that are found >80 times and with TF-target gene interactions that occur >80 times are considered. To avoid technical issues in the analysis, regulons with fewer than 8 target genes are removed from the final list. Subsequent analysis in Step 3 involves a stochastic downsampling to speed up computation, hence we verified that the chosen sample size was sufficient for accurate AUC approximations. We calculated n = 25 AUC matrices and confirmed that they contained few zeros and the variance of each matrix entry (i.e. approximated regulon activity in a given cell) was low.

### Bulk RNA deep sequencing analysis

The ‘Trim Galore!’ utility version 0.4.2 was used to remove sequencing adaptors and to quality trim individual reads with the q-parameter set to 20. Sequencing reads were then aligned to the mouse genome and transcriptome (Ensembl GRCm38 release-89) using RSEM version 1.3.0 (Li and Dewey, 2011) in conjunction with the STAR aligner version 2.5.2 (Dobin et al., 2013) . Sequencing quality of individual samples was assessed using FASTQC version 0.11.5 and RNA-SeQC version 1.1.8 (DeLuca et al., 2012). Differential gene expression was determined using the R-bioconductor package DESeq2 version 1.24.0 (Love et al., 2014). Gene set enrichment analysis (GSEA) was conducted as described in (Subramanian et al., 2005).

### Integration of single nucleus and bulk RNA deep sequencing data

Nuclei were simulated from each bulk RNA-seq replicate using a weighted random sampling of regulons with replacement. A regulon’s relative weight corresponded to the prevalence of its transcription factor in the given bulk RNA-seq sample. Each time a regulon was selected, the counts for its transcription factor and all target genes increased by one. The number of regulons expressed in each simulated nucleus was randomly selected from a list of how many unique transcription factors each snRNA-seq nucleus expressed (normalized expression > 2). Simulated nuclei were treated as raw count matrices and integrated with snRNA-seq nuclei using the SCT package in R (Hafemeister and Satija, 2019). To filter simulated nuclei, we trained 1000 linear discriminant analysis (LDA) classifiers with the python package scikit-learn (Pedregosa et al., 2011). For each classifier, snRNA-seq nuclei were split into test and train datasets, with 75% of nuclei used for training and the other 25% used for test. Each classifier was trained to predict whether a nucleus was a projection neuron (whether it was selected for subclustering in the initial Seurat analysis) based on values for the 30 top principal components from the SCT integration. Each classifier was applied to the remaining snRNA-seq nuclei for testing, and accuracy was evaluated using the Jaccard index calculated by scikit-learn (Pedregosa et al., 2011). The classifiers were then applied to the simulated nuclei. Simulated nuclei predicted to be projection neurons by all 1000 classifiers were designated as putative simulated projection neurons and selected for further analysis. Similarly, these putative simulated projection neurons were integrated with snRNA-seq mitral cells using SCT. 1000 LDA classifiers were trained to classify simulated nuclei as AON-projecting or PCx-projecting based on values for the 30 top principal components from the SCT integration. Each classifier was trained on 75% of the simulated projection neurons and tested on the other 25%, with accuracy evaluated using the Jaccard index. Each classifier was then applied to snRNA-seq mitral cells.

### smFISH in tracing experiments

Experiments were performed according to the manufacturer’s instructions, using the RNAscope Fluorescent Multiplex kit (Advanced Cell Diagnostics (ACD)) for fresh frozen tissue. Briefly, a total of 6 mice were injected into the AON and PCx with the rAAVretro-CAG-H2B-GFP. After 15 days post-injection, mice were deeply anaesthetized with 2.5% of 250mg/kg Avertin and transcardially perfused with 10 ml of ice-cold phosphate-buffered saline (PBS). The brains were dissected out from the skull, immediately embedded in Tissue Plus O.C.T compound (Fisher Healthcare) and snap frozen in a bath of 2-methylbutane on dry ice. Brains were cryo-sectioned coronally at 20 µm thickness, mounted on Fisherbrand™ Superfrost™ Plus microscope slides (Fisher Scientific) and stored at −80°C until use. *In situ* probes against the following mouse genes were ordered from ACD and multiplexed in the same permutations across sections: *Foxo1* (#485761-C2 and 485761), *Kcng1* (#514181-C2), *Lxh1* (#488581), *Sertm1* (#505401-C2), *Ebf3*(#576871-C3), *Sgcg* (#488051-C3), *Cadps2* (#529361-C3 and 529361), *Coch* (#530911-C3), *Ly6g6e* (#506391-C2), *Wnt5b* (#405051), *Fst* (#454331), *Barhl2* (#492331-C2), *Vdr* (524511-C3), *Gfp* (#409011, #409011-C2 and #409011-C3), *Piezo2* (#500501), *Olfr110/111* (#590641), *Calca* (#578771), *Lhx5* (#885621-C3), and *Vgll2* (#885631-C2). Following smFISH, high resolution images of a single z-plane were obtained using a 60x oil immersion objective on an Olympus FV3000 confocal microscope and a 40x oil immersion objective on a Nikon A1R-HD confocal microscope.

### Data and code availability

Raw single nucleus RNA and bulk RNA deep sequencing data have been deposited in Gene Expression Omnibus (GEO) under the accession numbers GSE162654 and GSE162655 respectively. The R and Python analysis scripts developed for this paper are available at the GitLab links https://gitlab.com/fleischmann-lab/molecular-characterization-of-projection-neuron-subtypes-in-the-mouse-olfactory-bulb and https://gitlab.inria.fr/acrombac/projection-neurons-mouse-olfactory-bulb. Extensive computational tools for additional in-depth exploration of our data sets are available through our website: https://biologic.crick.ac.uk/OB_projection_neurons.

## Acknowledgements

We thank the Crick Advanced Sequencing Facility, especially Robert Goldstone and Amelia Edwards for their excellent support. We thank Debipriya Das and Ana Agua-Doce from the Crick Flow Cytometry Facility for technical assistance, and the Crick animal facility for animal care. We thank the members of the Crick Digital Development Team, particularly Amy Strange, Luke Nightingale, Jude Pinnock and Marc Pollitt for excellent technical support. We thank the Harvard Neurobiology Imaging Facility for consultation and RNAscope services that supported this work. This facility is supported in part by the Neural Imaging Center as part of an NINDS P30 Core Center grant #NS072030. We thank Keeley Baker, Gilad Barnea, Bob Datta, Kevin Franks and Paul Greer for critical comments on the manuscript. Work in the ATS lab was supported by the Francis Crick Institute, which receives its core funding from Cancer Research UK (FC001153), the UK Medical Research Council (FC001153), and the Wellcome Trust (FC001153); a Wellcome Trust Investigator grant to ATS (110174/Z/15/Z), and a DFG postdoctoral fellowship to TA. Work is the AF lab was supported by grants from the NIH (1U19NS112953-01, 1R01DC017437-03) and the Robert J and Nancy D Carney Institute for Brain Science.

---

**Figure 1-figure supplement 1:**
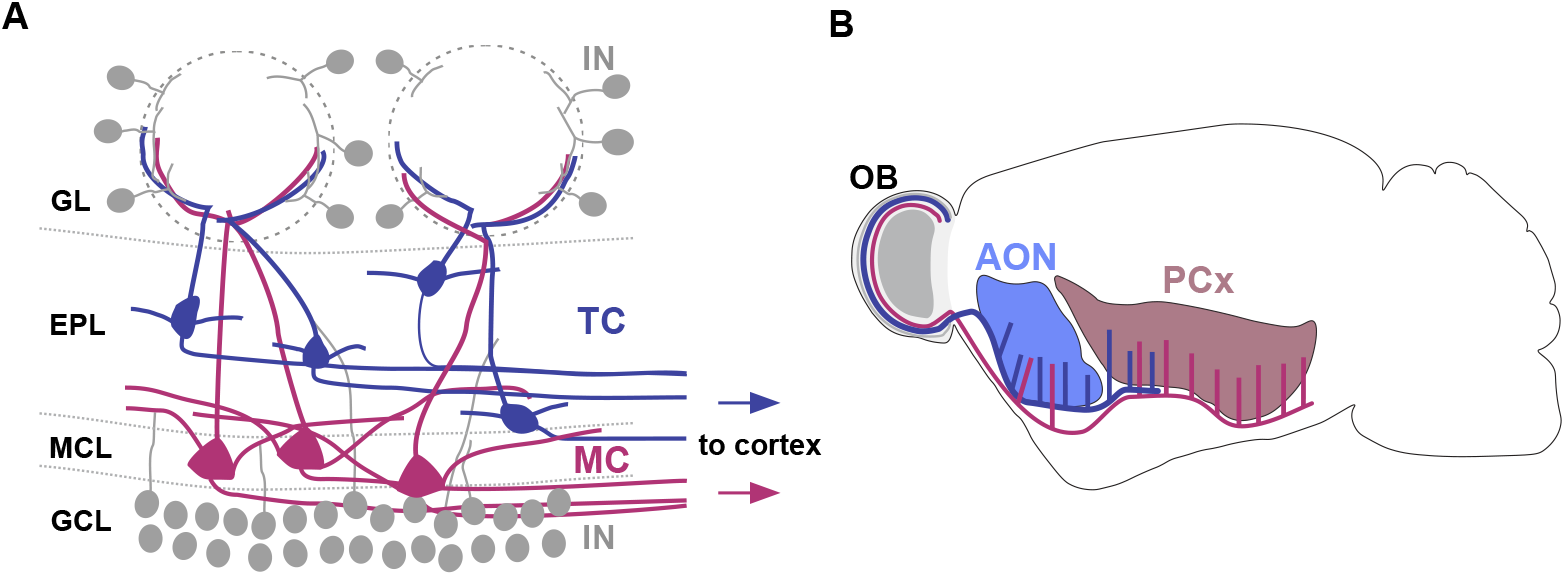
Schematic representation of olfactory bulb cell types and their cortical projection targets. **(A)** Schematic representation of cell types and their distribution within the olfactory bulb (IN: interneuron, TC: tufted cell, MC: mitral cell, GL: glomerular layer, EPL: external plexiform layer, MCL: mitral cell layer, GCL: granule cell layer). Tufted and mitral cells project their axons to downstream cortical regions. **(B)** Schematic representation of the distinct axonal projection targets for mitral and tufted cells in the olfactory cortex (AON: anterior olfactory nucleus, PCx: piriform cortex). Tufted cells project primarily to AON and to the anterior part of PCx, whereas mitral cell axons predominantly target the posterior portion of PCx.

**Figure 1-figure supplement 2:**
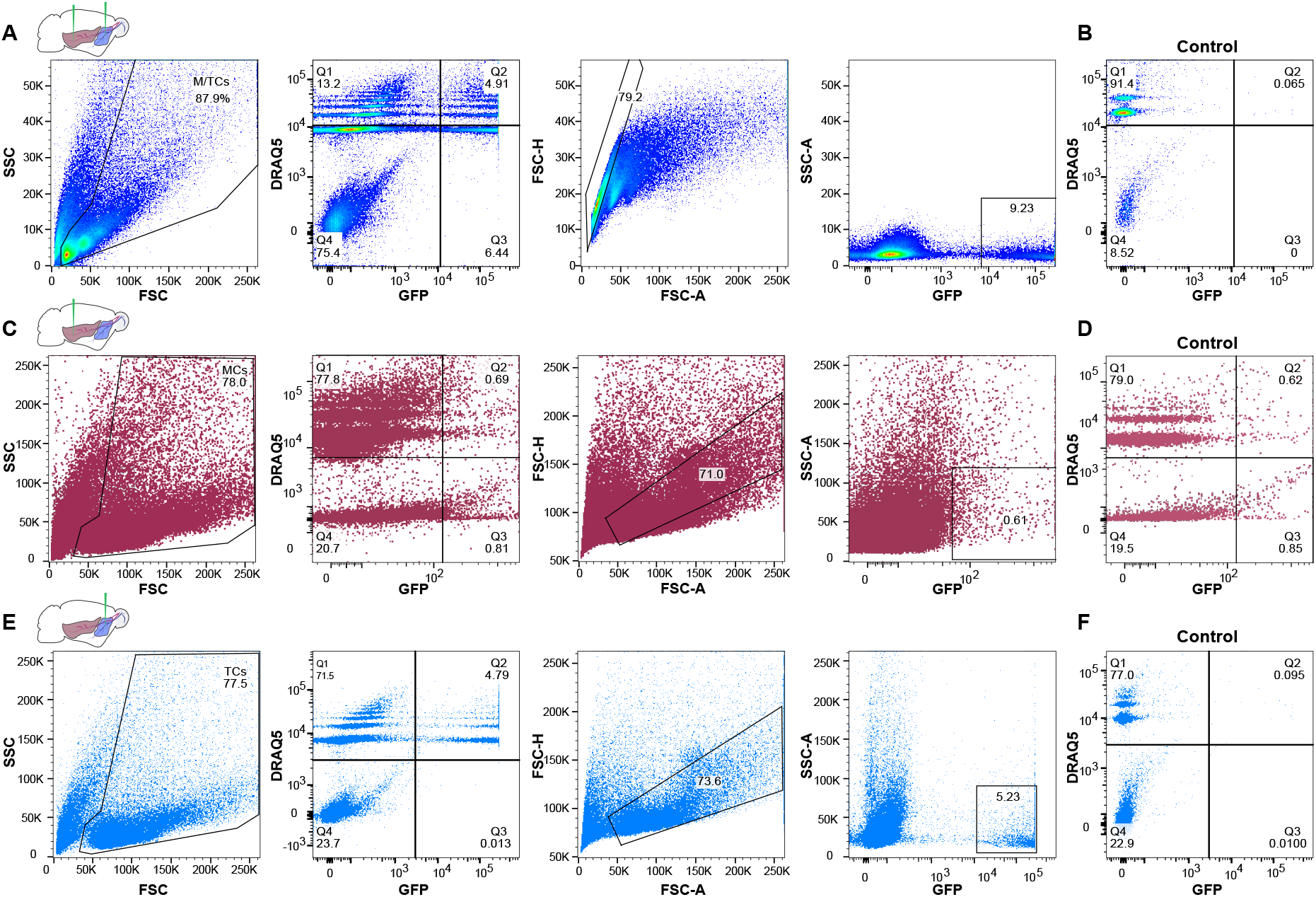
Enrichment of GFP-expressing nuclei using Fluorescence-Activated Nuclei Sorting (FANS). **(A)** Representative FANS data of GFP-expressing nuclei after injection of rAAVretro-CAG-H2B-GFP into AON and PCx to label OB projection neurons. From left to right: gating strategy for enrichment of GFP-expressing nuclei. Nuclei are identified based on size and granularity in the FSC (forward scatter) versus SSC (side scatter) plot. Next, the population of nuclei restricted to Quadrant 2 is selected that is both GFP-expressing and DRAQ5-positive (far-red fluorescent DNA dye). To exclude doublets, the population of nuclei around the diagonal in the FSC-A (forward scatter area) versus FSC-H (forward scatter height) plot is selected. Lastly, GFP-expressing nuclei filtered through the preceding steps are plotted against the SSC-A (side scatter area). **(B)** FANS results for control specimen for AON and PCx (olfactory bulb contralateral to the injection hemisphere from the same animal) showing the absence of GFP-expressing nuclei (same gating strategy as shown in **(A)**). **(C)** Representative FANS data after injection of rAAVretro-CAG-H2B-GFP into PCx to enrich for mitral cell nuclei. **(D)** FANS results for control specimen for PCx injections showing the absence of GFP-expressing nuclei (same gating strategy as shown in **(C)**). **(E)** Same as in **(C)** but after injection of rAAVretro-CAG-H2B-GFP into AON to enrich for tufted cell nuclei. **(F)** FANS results for control specimen for AON (olfactory bulb contralateral to the injection hemisphere from the same animal) showing the absence of GFP-expressing nuclei (same gating strategy as shown in **(E)**).

**Figure 2-figure supplement 1:**
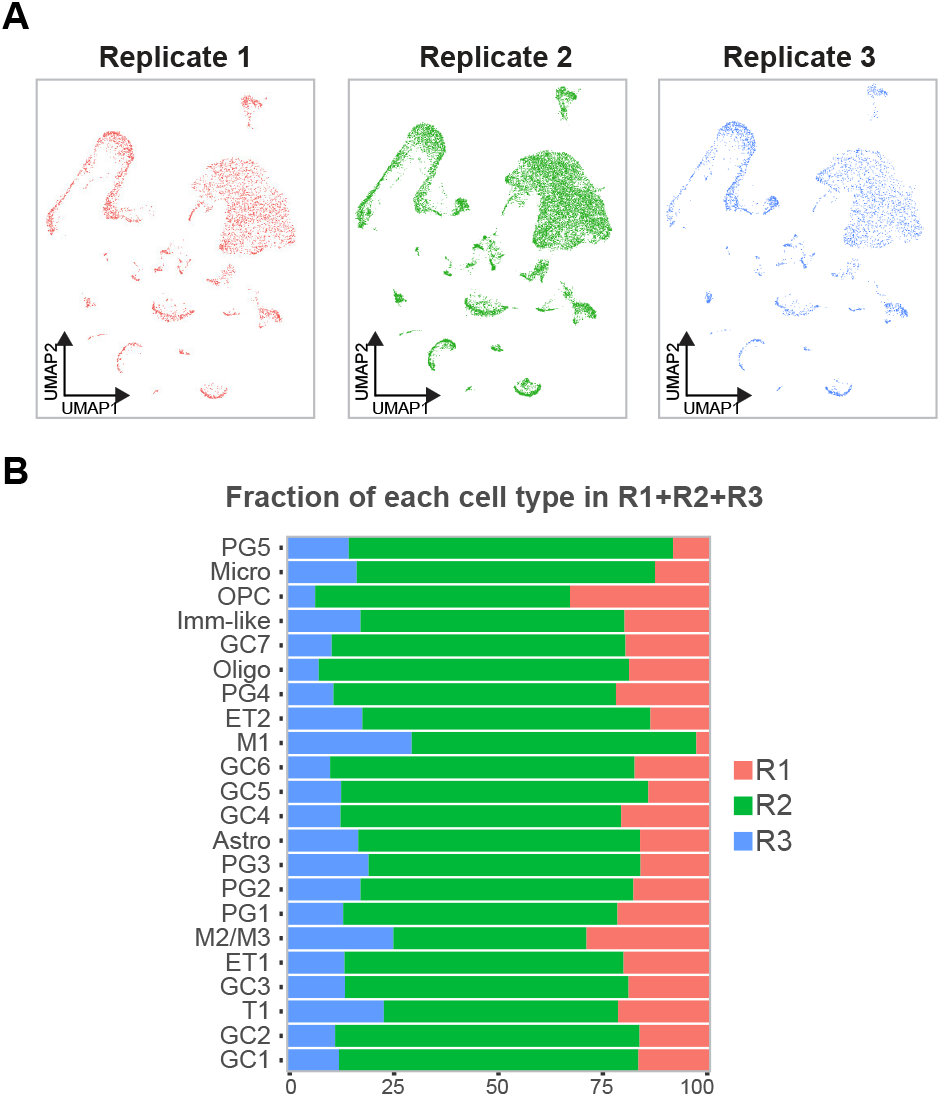
Quality check of individual replicates of sn-RNA-seq shows the reliability of the data and the replicability of each cell type. **(A)** Depiction of nuclei for each replicate (in red R1, in green R2, in blue R3) embedded in the UMAP space showing that replicates are very similar to each other and can be combined for downstream analyses. **(B)** Barchart showing the fraction of each cluster when combined R1, 2 and 3 color-coded by the replicate membership **(A)**, indicating that each cell type is consistently represented in each replicate.

**Figure 3-figure supplement 1:**
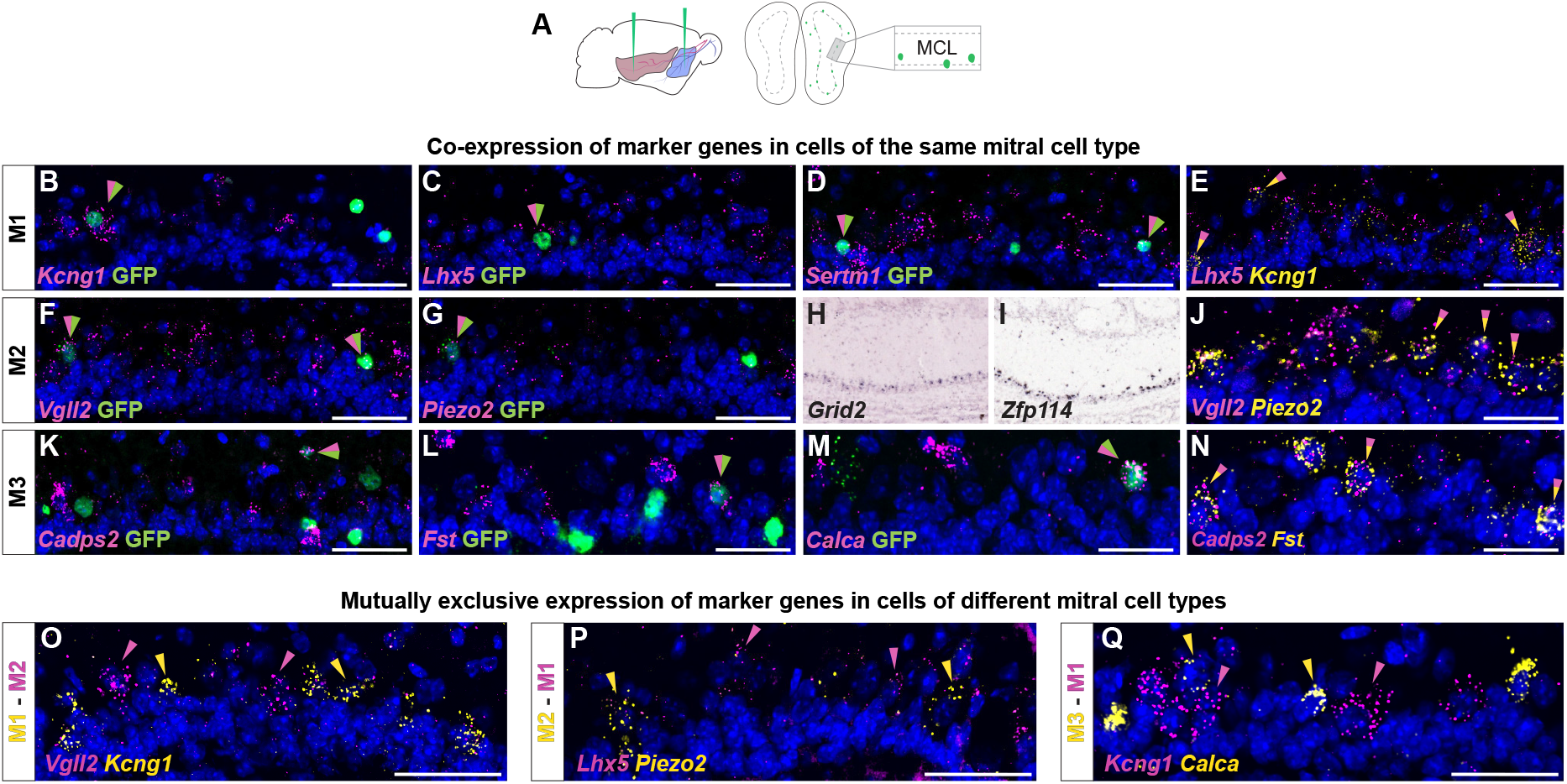
Histological analysis of DE genes for distinct mitral cell types. **(A)** Schematic representation of the smFISH images for validating selected mitral cell type-specific marker genes upon rAAVretro-CAG-H2B-GFP injection into PCx and AON. The scheme depicts the laminar location visualized in the histological images from a coronal section of the ipsilateral hemisphere to the injection site. MCL: mitral cell layer. **(B - D)** smFISH showing co-expression of M1-specific marker genes with viral GFP. Labeled nuclei are indicated by the magenta/green arrowheads. **(E)** smFISH showing co-expression of two M1-specific marker genes. Labeled nuclei are indicated by the yellow/magenta arrowheads. **(J)** smFISH showing co-expression of two M2-specific marker genes. Labeled nuclei are indicated by the yellow/magenta arrowheads. **(N)** smFISH showing co-expression of two M3-specific marker genes. Labeled nuclei are indicated by the yellow/magenta arrowheads. **(F, G)** smFISH showing co-expression of M2-specific marker genes with viral GFP. Labeled nuclei are indicated by the magenta/green arrowheads. **(H, I)** *In situ* hybridization images from the Allen Brain Atlas showing additional M2-specific DE genes. **(K - M)** smFISH showing co-expression of M3-specific marker genes with viral GFP. Labeled nuclei are indicated by the magenta/green arrowheads. **(O-P-Q)** smFISH images showing mutually exclusive expression of mitral cell type-specific marker genes for M1, M2, and M3. Yellow and magenta arrowheads show mutually exclusive expression of M1 versus M2 **(O-P)**, and M1 versus M3 **(Q)** marker genes. DAPI counterstain in blue. Scale bars, 50μm.

**Figure 3-figure supplement 2:**
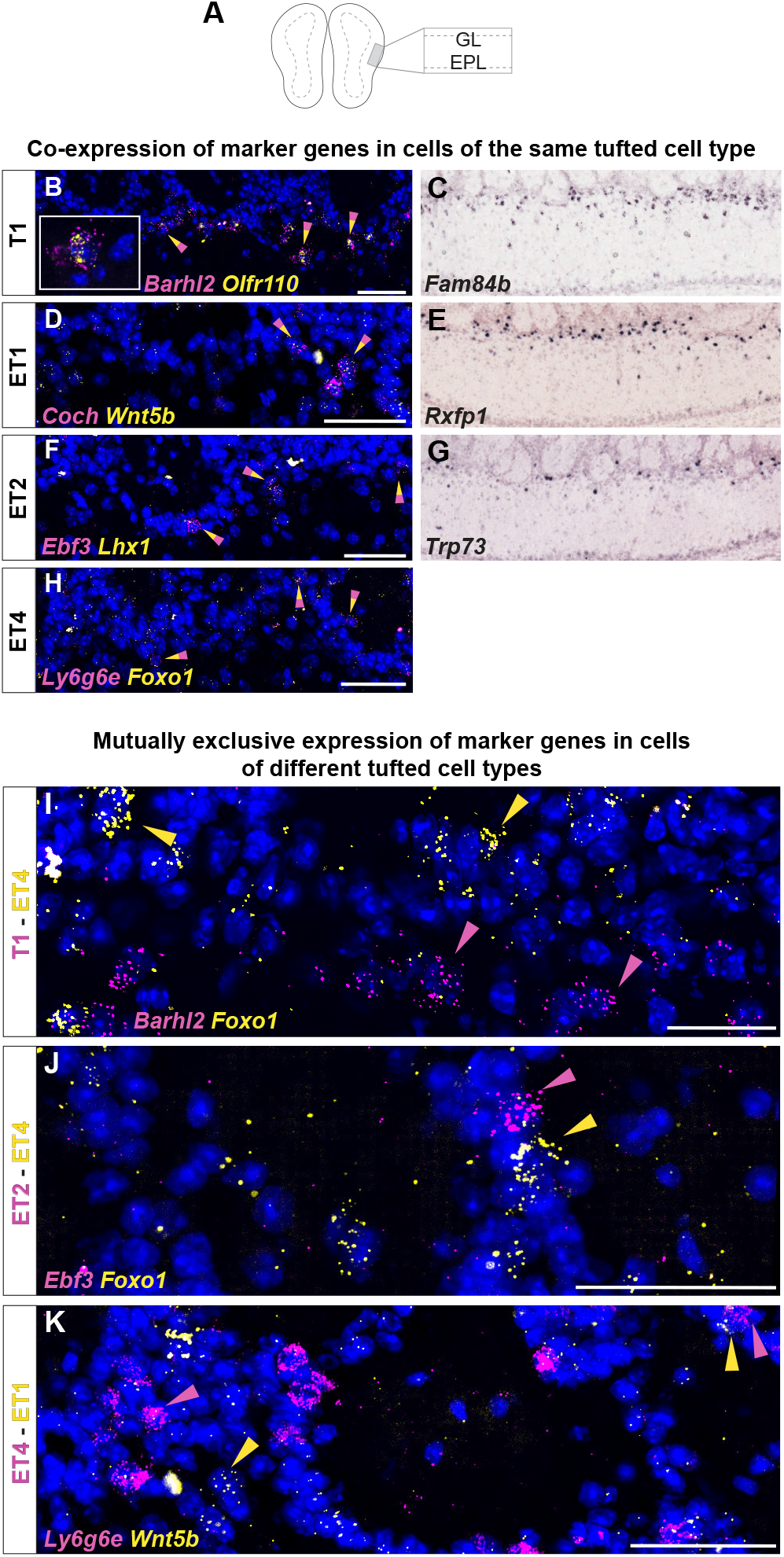
Histological analysis of DE genes for distinct tufted cell types. **(A)** Schematic of the smFISH images for validating selected tufted cell type-specific marker genes upon rAAVretro-CAG-H2B-GFP injection into PCx and AON. The scheme depicts the laminar location visualized in the histological images from a coronal section of the ipsilateral hemisphere to the injection site. GL=glomerular layer; EPL=external plexiform layer. **(B)** smFISH showing co-expression of two T1-specific marker genes. Labeled nuclei are indicated by the yellow/magenta arrowheads. High magnification (left) shows clear co-labeling of the two mRNA probes *Barhl2* and *Olfr110*. **(C)** *In situ* hybridization images from the Allen Brain Atlas showing one additional T1-specific DE gene. **(D)** smFISH showing co-expression of two ET1-specific marker genes. Labeled nuclei are indicated by the yellow/magenta arrowheads. **(E)** *In situ* hybridization images from the Allen Brain Atlas showing additional ET1-specific DE genes. **(F)** smFISH showing co-expression of two ET2-specific marker genes. Labeled nuclei are indicated by the yellow/magenta arrowheads. **(G)** *In situ* hybridization images from the Allen Brain Atlas showing additional ET2-specific DE genes. **(H)** smFISH showing co-expression of two ET4-specific marker genes. Labeled nuclei are indicated by the yellow/magenta arrowheads. **(I-J-K)** smFISH images showing mutually exclusive expression of tufted cell type-specific marker genes for T1, ET1, ET2 and ET4. Yellow and magenta arrowheads show mutually exclusive expression of T1 versus ET4 **(I)**, ET2 versusET4 **(J)**, and ET1 versus ET4 **(K)** marker genes. DAPI counterstain in blue. Scale bars, 50μm.

**Figure 4-figure supplement 1:**
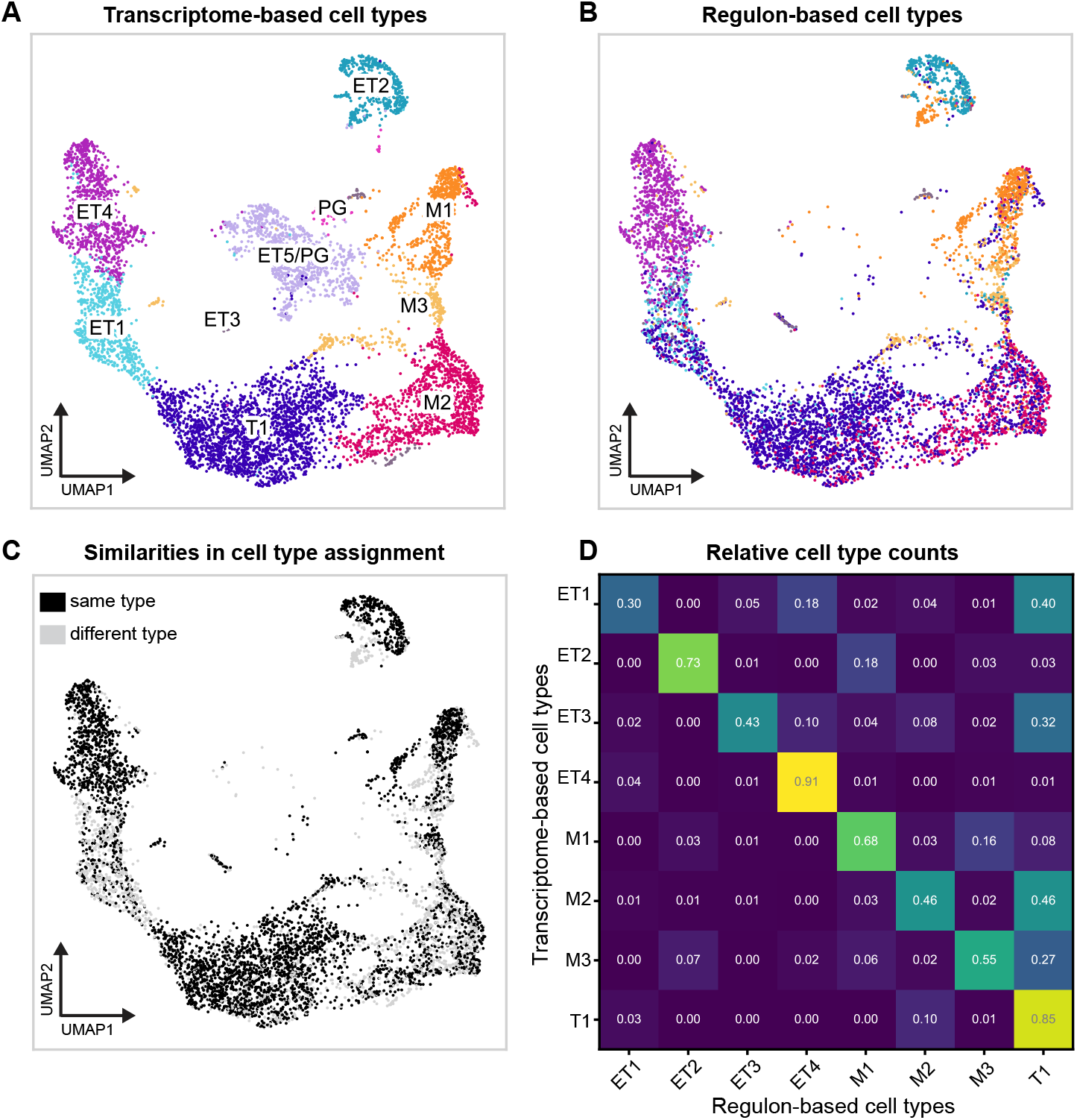
Regulon-based clustering and transcriptome-based clustering provide complementary axes for cell type identification. **(A)** Leiden clustering of projection neurons in a UMAP space computed from regulon activity, visualized on the UMAP coordinates computed from the whole transcriptome. The major cell type clusters are consistent between the two approaches, even if boundaries are more diffuse and have shifted. Regulon-based clusters were computed through a fine-grained clustering of cells on their regulon activity score. We computed a knn graph (nr of neighbors=10) and then clustered cells with the Leiden algorithm (resolution = 5.5), resulting in 86 clusters. Regulon-based cell types were named for the most abundant transcriptome-based cell type present in the given cluster. This reduced the 86 clusters to 8. Note that the Leiden algorithm is an improved version of the Louvain clustering algorithm (Traag 2018, arxiv). **(B)** UMAP of cells assigned the same type (black) or a different type (grey) by both methods. **(C)** Overlap between transcriptome-based and regulon-based cell type assignment. Each entry represents the fraction of cells in a given transcriptome-based cell type that were assigned to a given regulon-based cell type. Cell type assignments were mainly consistent, with changes mostly observed in the expansion of T1 under regulon-based clustering to include many cells from ET1, ET3, M2 and M3.

**Figure 4-figure supplement 2:**
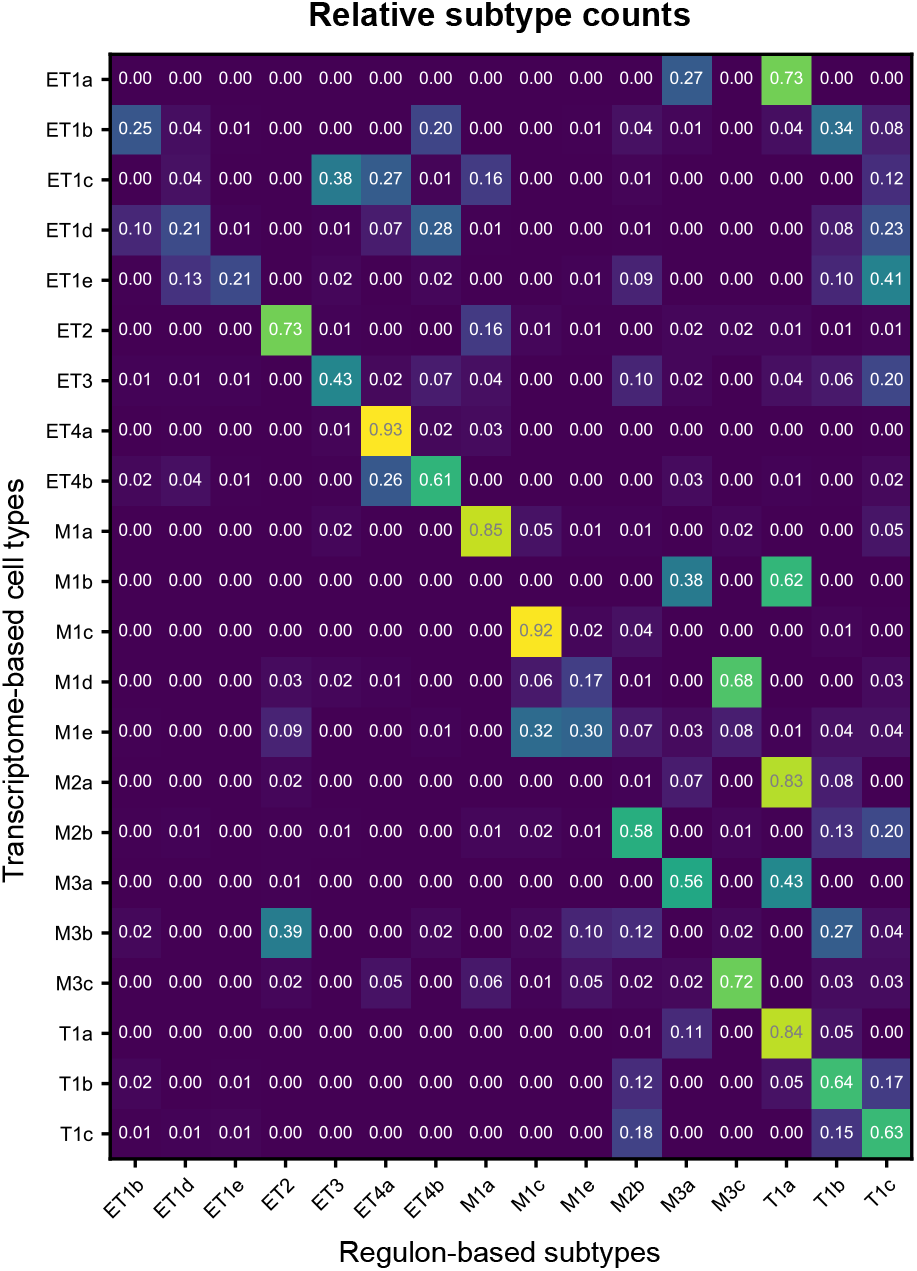
Overlap between transcriptome-based and regulon-based cell subtype assignment. Subtypes were assigned using the method described in **Figure 4-figure supplement 1A**. Each entry represents the fraction of cells in a given transcriptome-based cell subtype that were assigned to a given regulon-based cell subtype. Differences are mainly observed in the expansion of T1a, M3c and ET2 in regulon-based assignment.

